# Impact of Infectious Diseases on Wild Bovidae Populations in Thailand: Insights from Population Modelling and Disease Dynamics

**DOI:** 10.1101/2023.08.29.554960

**Authors:** Wantida Horpiencharoen, Jonathan C. Marshall, Renata L. Muylaert, Reju Sam John, David T. S. Hayman

## Abstract

The wildlife and livestock interface is vital for wildlife conservation and habitat management. Infectious diseases maintained by domestic species may impact threatened species such as Asian bovids, as they share natural resources and habitats. To predict the population impact of infectious diseases with different traits, we used stochastic mathematical models to simulate the population dynamics 100 times over 100 years for a model gaur (*Bos gaurus*) population with and without disease. We simulated repeated introductions from a reservoir, such as domestic cattle. We selected six bovine infectious diseases; anthrax, bovine tuberculosis, hemorrhagic septicaemia, lumpy skin disease, foot and mouth disease and brucellosis, all of which have caused outbreaks in wildlife populations. From a starting population of 300, the disease-free population increased by an average of 228% over 100 years. Brucellosis with frequency-dependent transmission showed the highest average population declines (−97%), with population extinction occurring 16% of the time. Foot and mouth disease with frequency-dependent transmission showed the lowest impact, with an average population increase of 200%. Overall, acute infections with very high or low fatality had the lowest impact, whereas chronic infections produced the greatest population decline. These results may help disease management and surveillance strategies support wildlife conservation.

## Introduction

Livestock encroachment into wildlife habitats can drive disease transmission between wildlife and domestic livestock, which is a vital issue for both human public health and wildlife conservation. An effect of agricultural expansion and land-use change is to bring wildlife and livestock close to each other and increase the contact frequency and time between domestic and wildlife populations (1–3). This increased contact may increase the risk of disease transmission as they can share the same natural resources (e.g. grassland and water) (4).

Infectious diseases can cause dramatic declines in wildlife populations, as demonstrated by chytridiomycosis, which has been implicated in the likely extinction of over 200 amphibian species (5). Most infectious bovid pathogens are capable of infecting both domestic and wild species. For example, bighorn sheep populations declined from ovine respiratory disease (*Mycoplasma ovipneumoniae*) acquired when sharing the grazing areas with domestic sheep (6). Similarly, bovine brucellosis has been transmitted from domesticated yak to wild yak in China (7), and between bison, elk and domestic cattle in the USA (8). Brucellosis affects these bison and elk populations both indirectly and directly as the seropositive animals may be culled for management and directly as the pathogen affects animal reproductive systems (9). Critically, the impact of infectious diseases is determined by disease-specific traits, such as infection fatality rates (10).

There are five wild bovids species (gaur, banteng, wild water buffalo, mainland serow and Chinese goral) that remain in Thailand. They are experiencing dramatic population declines from habitat destruction, illegal hunting (11), and resource competition with domestic livestock (12). Infectious diseases transmitted from contact with domestic cattle could cause further declines. Several diseases circulate in Thai cattle, including endemic diseases like bovine tuberculosis (bTB) from *Mycobacterium bovis* (13), and new infectious diseases, such as the recent lumpy skin disease (LSD) (14).

Infectious disease modelling provides a tool to understand disease dynamics better and predict the potential consequences of infection in a population, helping disease prevention and control programs (15), particularly as collecting field data or conducting experiments on some pathogens and hosts is extremely challenging. Models have, for example, been used to determine the potential impact of disease on endangered species, such as canine distemper in the Amur tiger (16). Although models contain uncertainty and may not cover all factors, predictions can guide the policies and help decision-making (17).

Here, we use mathematical models to explore the potential consequences of six major bovine infectious diseases on endangered Thai wild bovid populations. Our aim is to estimate the potential population changes after the disease is introduced in the population from a reservoir, such as domestic cattle. The diseases are anthrax, haemorrhagic septicaemia (HS), bTB, LSD, foot and mouth disease (FMD) and bovine brucellosis, which all infect a range of bovid species, are distributed worldwide, including Thailand, and have different characteristics. Our study predominantly focuses on the gaur (*Bos gaurus*) population as their populations are well described plus, of five species of Thai wild bovids, they have the greatest opportunity to interact with domestic livestock and humans since they are the most likely to share space and resources (e.g. agricultural areas, watering holes) (18, 19). We hypothesised that acute infections with very low and very high infection fatality rates would have less impact on populations than those with moderate mortality or chronic diseases, the latter high fatality case because they ‘burn out’ by removing infectious individuals rapidly (10). The study aims to help infectious disease surveillance and monitoring prioritisation strategies in wildlife and livestock for wild bovid conservation.

## Material and methods

### 1. Model construction

#### 1.1. Population dynamic models

We selected gaur populations as a model system because they are widespread across Thailand, overlap with livestock and people, and demographic data are available (18, 20). Further, their demography is similar to other threatened wild bovids (Figure S1 in supplementary material). We used the same model structure for all species. The demographic parameters for the remaining four bovid species used in simulations are provided in Table 2 because they exhibit variations in population sizes, social behaviours, and distribution, making them interesting for further infectious disease modelling of population impact.

**Table 1.** Diseases, pathogens, and the structures adopted in the modelling procedures.

| Disease | Pathogens | Model structure |
| --- | --- | --- |
| Anthrax | *Bacillus anthracis* | $S \rightarrow I$ |
| Bovine TB | *Mycobacterium tuberculosis* | $S \rightarrow E \rightarrow I$ |
| Haemorrhagic septicaemia | *Pasteurella multocida* | $S \rightarrow I \rightarrow R \rightarrow S$ |
| Lumpy Skin Disease | *Capripoxvirus* | $S \rightarrow E \rightarrow I \rightarrow R \rightarrow S$ |
| Foot and Mouth Disease | *Aphthovirus* | $S \rightarrow E \rightarrow I \rightarrow R \rightarrow M \rightarrow S/E$ |
| Bovine brucellosis | *Brucella abortus* | $S \rightarrow E \rightarrow I \rightarrow R \rightarrow M \rightarrow S/E$ |

**Table 2.**
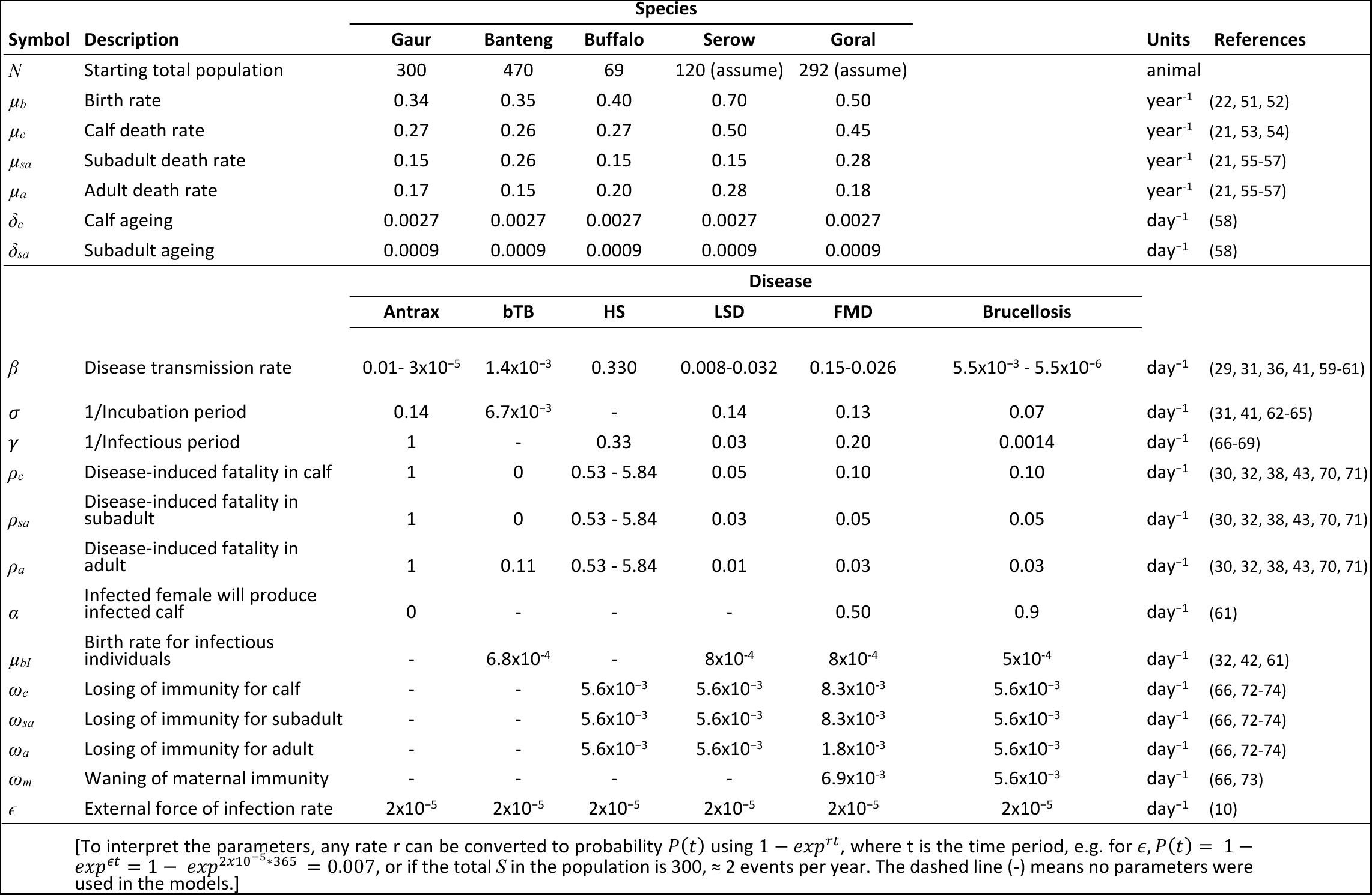
Parameters and variables.

We assumed demographic parameters were otherwise constant. If *N* is the total animal population, *N_a_* is the adult population, *N_sa_* is the subadult population, *N_c_* is the calf population and *µ* is the annual birth rate. Only adult females were assumed to add new calves to the population, which enter the susceptible class at a birth rate *µ_b_N_a_*. Animals can leave their compartments at the natural death rate (*µ_a_, µ_sa_* or *µ_c_*) or age from calf to subadult (*δ_c_*) and from subadult to adult (*δ_sa_*). The natural death rate was estimated based on the mortality rate of wild ungulates and gaur in captivity (21). The initial population was 300 animals, based on the gaur population size in the Khao Pheang Ma non-hunting area (8 km^2^) in Thailand (22, 23). Thus, the population dynamic model equations at time *t* can be as following:

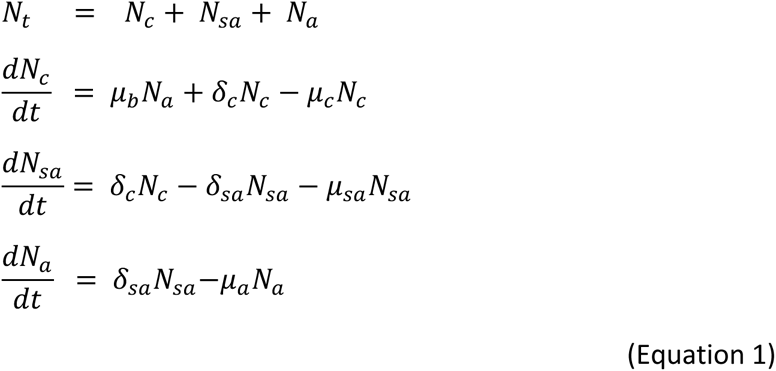

#### 1.2. Infectious disease models

We used the same age-structured population as the baseline model (Equation 1) and incorporated compartments with different parameter values for building the disease models.

We modelled the diseases based on Susceptible-Infected-Recovered (*SIR*) models and modified them based on the disease parameters of domestic animals (e.g. dairy cattle, domesticated buffalo) and wildlife from previous studies and background knowledge. Table 1 presents the diseases and model structures we used, and a flow diagram is in the supplementary materials. For the compartments used in the models, *S* denotes the number of susceptible animals, *E* denotes the number of exposed animals, *I* denotes the number of infected animals, *R* denotes the number of recovered animals, and *M* denotes the number of calves with maternally derived immunity.

We selected six infectious diseases which have been reported to cause outbreaks in wild ungulates and livestock populations in several places, including Thailand, which are: anthrax (*Bacillus anthracis*) with an *SI* structure, bovine tuberculosis (bTB-*Mycobacterium bovis*) with an *SEI* structure, haemorrhagic septicaemia (HS-*Pasteurella multocida*) with an *SIRS* structure, lumpy skin disease (LSD) with an *SEIRS* structure, and both foot and mouth disease (FMD) and brucellosis (*Brucella abortus*) with an *SIERMS/E* structure. Table 2 displays the disease parameters used in the models. These infections have a range of key parameters of interest. They include infectious diseases with very short (effectively no) incubation periods (e.g. HS) to long incubation periods (e.g. bTB), and very high mortality (e.g. anthrax) to low mortality (e.g. LSD, FMD).

#### 1.3. Mode of transmission

Different transmission types can provide different model results (24). Here, we considered two disease transmission modes; 1) density-dependent (DD) and 2) frequency-dependent (FD). DD transmission is assumed when the contact rate is proportionate to the population density, while FD transmission is assumed when the contact rate is independent of the population density (24, 25). We assumed the transmission modes for each pathogen and then compared them by introducing both transmission modes because, for some infections, there is no clear evidence of which type suits the pathogen’s transmissions and these represent extreme situations of population change for both transmission modes (10).

Transmission is often likely a mix of both DD and FD in many cases such as FMD and bTB (26, 27). For the transmission rate (*β*), we used parameter values based on the reference studies with the reported FD or DD transmission (Table 2), which differs among infectious diseases. However, to test the sensitivity of the results to these assumptions, we also rescaled the *β* rate to all models to examine the consistency of the results between FD and DD using Equations 2:

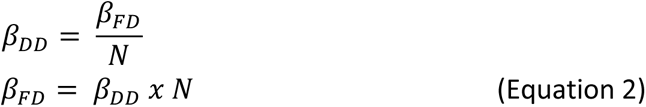

#### 1.4. Infection reintroduction

To model the repeated introduction of an infection from a reservoir such as domestic cattle (e.g. for FMD) or the environment (e.g. anthrax), we repeatedly reintroduced infection into our population at rate *ɛ* independently of any infection in the population. This reintroduction means the impact of infections are not simply estimated by the basic reproductive number (*R*_0_), the average number of secondary cases caused by a primary case in a completely susceptible population.

### Anthrax (Bacillus anthracis)

To model anthrax, we initially assumed the transmission is FD. We used an *SI* model (28, 29) with the transmission rate for FD at 0.01, then rescaled to FD using Equation 2. We assumed that *S* animals are exposed to infected animals and then become infectious (*I*) at rate *β*. All infected animals (*I*) die (100% mortality) (30) at disease-induced death rate (*ρ*) and the infectious rate (*γ*), *γρI*.

### Bovine tuberculosis (bTB-*Mycobacterium tuberculosis*)

Bovine TB is a chronic and zoonotic infection in livestock and wildlife worldwide (31). We first assumed DD transmission and used an *SEI* structure for modelling. The flow of the model starts from *S*, which are exposed to *I* animals and become exposed (*E*) at transmission rate (*β*); then *E* animals enter the *I* compartment at the incubation rate (*σ*). As we assume lifelong infection without recovery (31); *I* animals either die with an age-specific disease-induced fatality rate (*ρ*) or natural death rate (*µ*). *S* and *E* adults give birth with the normal birth rate *µ_b_*(*S_a_* + *E_a_*) but *I* adults are assumed to have a lower fecundity rate (reduced by 27%, (32) at *µ_bI_I_a_*. Bovine TB has a long incubation period from several months up to 7 years (33), so here we used 5 months based on the mean incubation period in the African buffalo (31). We also assumed there’s no vertical or pseudo-vertical (e.g. in utero or calf rearing) transmission as it is uncommon for bTb (34, 35).

### Haemorrhagic septicaemia (HS-*Pasteurella multocida*)

HS is a fatal septicaemic disease in cattle and buffalo. We assume DD transmission based on a previous HS modelling study (36). We used an *SIRS* model and excluded an *E* class as the disease can show acute clinical signs with a short incubation period of ∼ 18-20 hours (37) animals become *I* at the transmission rate (*β*). *I* animals may die from HS at the disease-induced fatality rate (*ρ*) or survive and become recovered (*R*) at infectious rate (1*/γ*). We calculated the fatality rate in the cattle population to range from 0.53% to 5.84%. This was determined by dividing the number of deaths from HS (0.21%, assumed from the percentage of deaths from bovine respiratory disease, (38) by the minimum (3.59%) and maximum (40%) prevalence of seropositive animals from *P. multocida* infected herds (39, 40). Therefore, we used two infection fatality rates (0.53 and 5.83%), since case fatality is underestimated, as a large proportion of animals are infected but do not develop clinical signs of diseases. *R* animals reenter *S* when they lose immunity at the immunity loss rate (*ω*). We used the proportion of susceptible animals (0.6) to calculate *R*_0_ and therefore the *β* rate using the equation: *R*_0_ = (1*/* (1− *I*)) = 1*/S*.

### Lumpy skin disease (LSD-Lumpy skin disease virus, *Capripoxvirus*)

We used an *SEIRS* structure for LSD. We inserted *E* and *R* compartments as the disease has an incubation period of between 7 and 14 days and a recovery period of around 4-6 months. We initially assumed the transmission is DD as the cattle density could be one of the risk factors to increase the transmission rate within-herd. However, we used both FD and DD *β* values because the published work has reported differences in incidence rates associated with different transmission modes (41). We assumed different birth rates for *I* females (*µ_bI_*) from the natural birth rate, because LSD can reduce the fertility rate by 10% (42). Also, we applied the highest fatality rate in calves (5%) and lower mortality rates to subadults (3%) and adults (1%) (43).

### Foot and mouth disease (FMD-Foot-and-mouth disease virus, *Aphthovirus*) & Bovine Brucellosis (*Brucella abortus*)

We initially assumed the transmission was FD for both FMD and brucellosis. An *SEIRMS/E* model was applied for FMD and brucellosis. We considered the *SEIR* model appropriate for both diseases. Recovered FMD and brucellosis cows can pass immunity to their offspring. Therefore, we added a maternally-derived immunity (*M*) compartment, which refers to the calves born with maternally-derived immunity from recovered mothers (*R_a_*). We assumed that if recovered adults (*R_a_*) calve at the birth rate (*µ_b_*), a calf will receive maternal immunity and stay in *M* compartment for an average of 6 months (44) before immunity wanes and they become susceptible again (*S_m_*) at a loss of immunity rate (*ω_m_*). *S_m_* calves can either become an exposed calf (*E_c_*) if there is contact with *I* or enter a susceptible subadult (*S_sa_*) compartment at a loss of immunity rate plus calf ageing rate: 1/*δ_c_* = 1/(*δ_m_* + *ω_m_*) or 1/*δ_m_* = 1/(*δ_c_*-*ω_m_*) if they have no contact with *I* to ensure that animals spend the same average time in the calf age class (*_c_*).

Vertical transmission from mothers to calves can be a consequence of infection among infected mothers with different probabilities for FMD (∼ 0.5) and brucellosis (∼ 0.9). Infectious adults are assumed to produce an infectious calf (*I_c_*) entering *I* at the birth rate *µ_b_I*. The proportion of infected females producing infected calf denotes *α*. So, an infected female can produce an infected calf at a rate *αµ_bI_I_a_* and produce a susceptible calf at a rate (1−*α*)*µ_bI_I_a_*.

The mathematical ordinary differential equations for the calf population (*X_c_*) are:

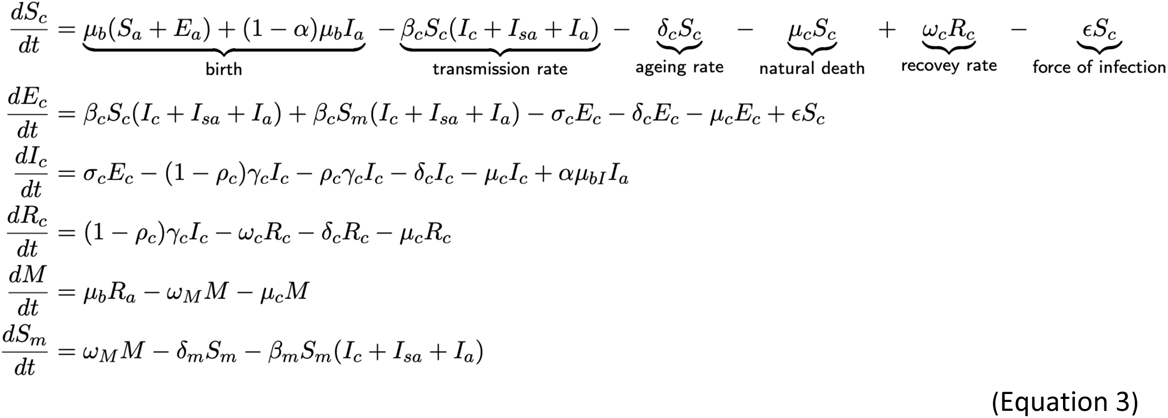

The system of ordinary differential equations and all other disease model equations and diagrams can be found in the supplementary materials. Note that all equations are subsets or variants of Equation 3.

### 2. Model simulations

Due to the small population size of gaur and other endangered bovids, we were interested in how infections might lead to their decline. Additionally, as we aimed to allow infections to go extinct in populations if they could not be sustained. Therefore, we chose stochastic models for this study because they effectively capture the stochastic nature of wildlife populations using random values. This randomness introduces variation in population sizes, which significantly affects small population sizes and long-term simulations (45). First, we built the population dynamics model without infectious disease classes and parameters as a baseline model (10). Then, we introduced an infectious adult (*I_a_* = 1) to the susceptible (*S*) population. We assumed that *I* would infect *S* at a transmission rate, *β*, and enter the next compartment based on the model structure. Demography (birth rate, natural death rate and ageing rate), the external force of infection (*ɛ*), and disease-induced fatality (*ρ*) were included in all disease models. The stochastic simulation was performed using the Poisson distribution to calculate the probability of events by multiplying the rate parameters *i* with a time step through Gillespie’s *τ*-leap algorithm (*τ* = 1) (Equation 4).

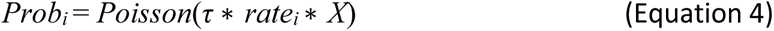

Where *X* is a state (e.g. *S, I, R*). All models were simulated for 100 years, and the stochastic models were simulated 100 times to generate the mean and to understand the uncertainty. We modelled the population change for 100 years, as long-term simulations of at least 10 years or three generations of species is recommended to explore population trends, and short-term time series may lead to misleading conclusions (46).

The parameter values used for modelling were collected from the literature review and observational data (Table 2).

### 3. Measuring impact

We compared the difference in total population (*N*) between no infection and disease models by calculating the average percentage of the population change using the total population at the start (*N_t_*_=0_) minus the total population at the end, (*N_t_*_=100_) of the simulation time, divided by *N_t_*_=0_ and converted this to a percentage, then divided by 100 times of simulations, using the following equation:

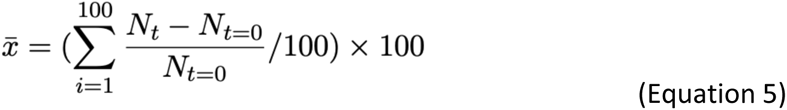

We used a principal component analysis (PCA) to find which diseases showed similar traits group by four disease parameters (transmission rate, incubation rate, infectious rate and fatality rate) which were included in all models and then coloured the values based on the percentage of the total population change. We performed PCA in R software using the PCATools package (47). The highest percentage of the first two axes contributed most to the population percentage changes.

### 4. Code availability

We used R Core Team (48) to simulate all the models and for further analysis. The R code for reproducing the analyses is available at a GitHub repository https://github.com/Wantidah/InfectiousModel.

## Results

### 1. Disease free model

We developed stochastic models for a gaur population, including a baseline model without infection and six infectious disease models. The baseline model of the gaur population demonstrated significant population growth, increasing from 300 to an average of 685 (range 113-1469) additional animals, which is approximately a 228% (38-489%) increase over the 100 years simulated. The average adult and subadult populations consistently increased, while the calf population slightly decreased from 95 to 82 animals (–279) on average (Figure 1: A&I). This gave us a disease-free population to model the impact of disease introduction into.

**Figure 1.**
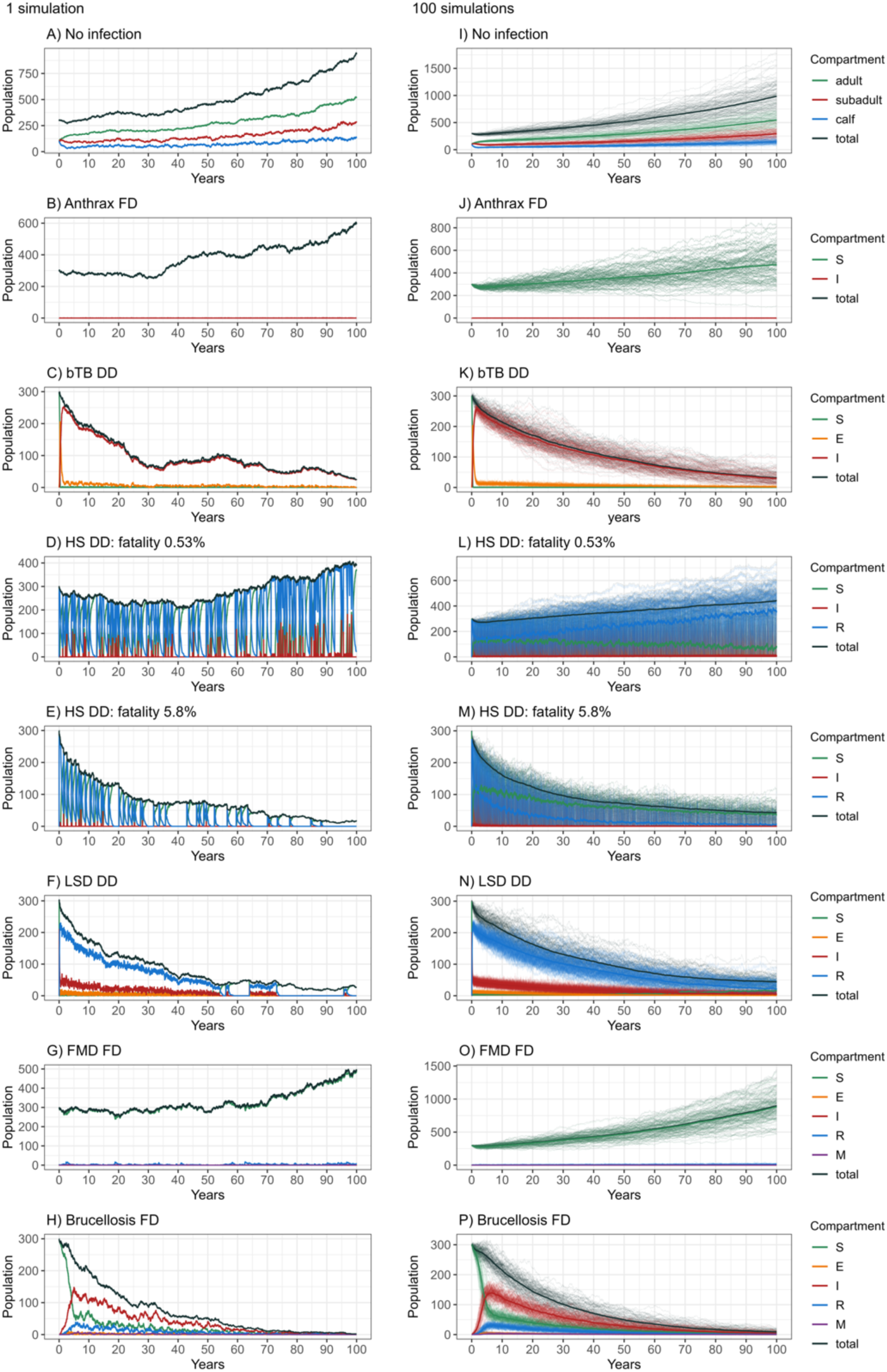
Modelled gaur population dynamics with and without disease: A-H are single example stochastic simulations for 100 years; I-P are 100 stochastic simulations for 100 years. Mean values are in solid lines. A and I are no infection models and the others are the infectious disease models where bTB is bovine tuberculosis; HS haemorrhagic septicaemia; LSD lumpy skin disease; and FMD foot and mouth disease. The entire model results, including all disease parameters used in the simulations, can be found in the supplementary materials.

The population dynamics of the four other wild bovid species in Thailand show similar trends to the gaur population (Figure S1, Figure 1), so we assumed there will be similar trends for the other two large bovids (banteng, wild water buffalo) that have similar herd sizes, population demography (e.g. age-structured, birth rate, death rate) and social behaviours to gaur (49).

However, the population dynamics may differ from the medium-sized bovids (Chinese goral and mainland serow) that live in smaller groups or even pairs and can be isolated from each other (50).

### 2. Disease impacts

Brucellosis had the greatest impact on population decline, while FMD had the lowest impact. Our PCA quantitatively shows that pathogens with longer incubation periods, chronic infection and medium to low fatality lead to greater population decline in smaller populations of endangered bovids than a high fatality or high transmission rate alone. Most diseases were grouped by similar traits which can see in the PCA biplot for brucellosis, bTB and LSD, while a few variations were seen for HS, and the FMD FD model was something of an outlier, (rescaling DD) at *β* = 6552 (Figure S17 and Figure S18). The greatest contribution to the percentage population change in the first axis was the infectious rate (55%) and fatality rate (42%). For the second axis, is the incubation period (74%). The first axis, PC1, has 43.25% and second axis PC2 has 31.61% of the variance explained.

Using different parameter values, fatality rates and modes of transmission yielded different effects on the modelled populations for HS, FMD, LSD and brucellosis. FD brucellosis had the largest population impact, yet DD brucellosis suppressed population growth but led to a stable population. In contrast, FD transmission of HS, LSD and FMD showed a continued population increase. Anthrax and bTB showed only a slight difference in the average population change between the two transmission modes (Figure 3; (A)). Simply rescaling the *β* with modelling FD or DD transmission had limited changes, which demonstrated consistency in the population change within the same infectious disease. Rescaling the *β* also reduced the probability of local extinction in the gaur population (*N* = 0) for FD brucellosis. Figure 2 and Figure 3 present the results of rescaling.

**Figure 2.**
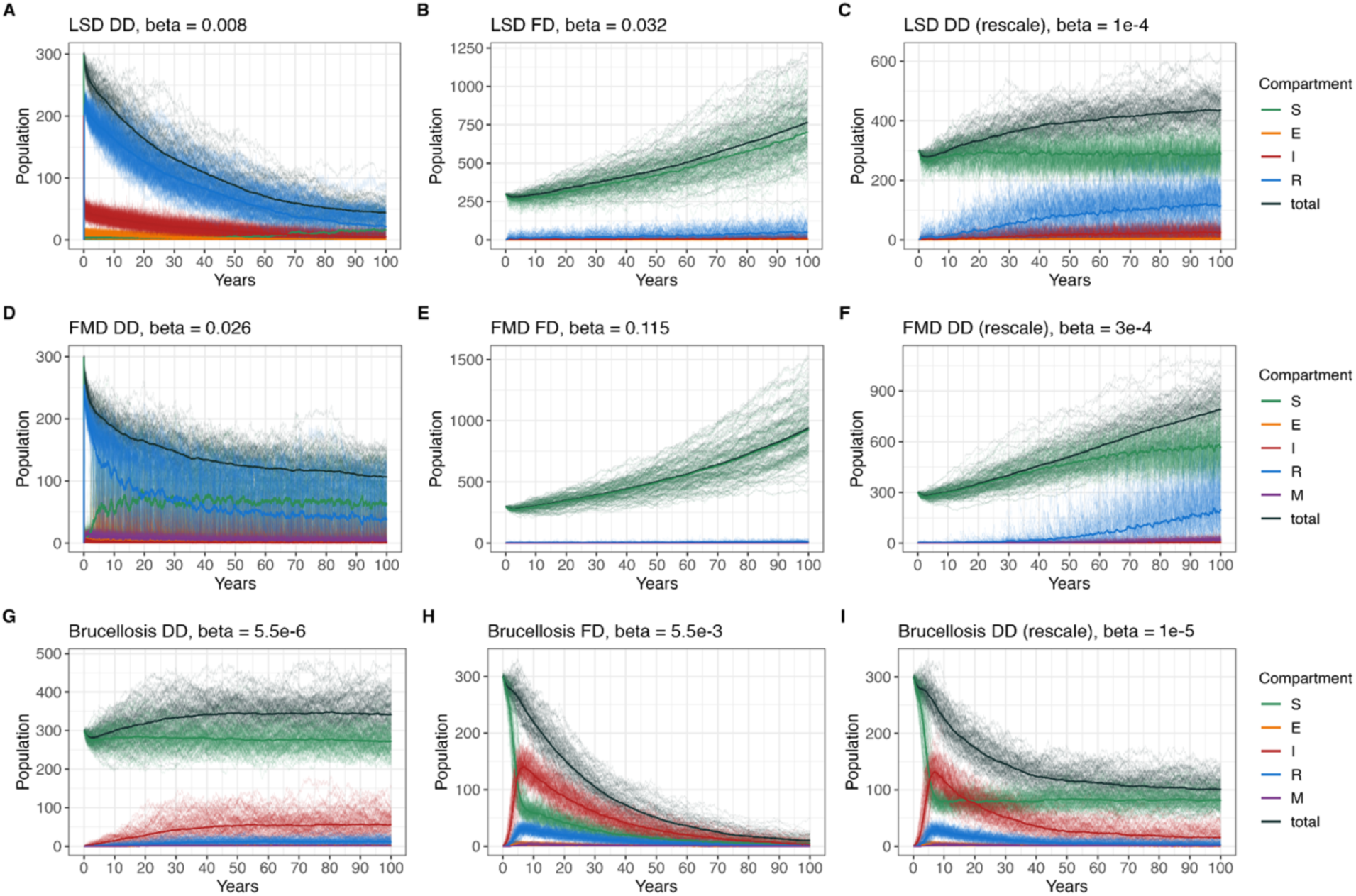
Population dynamics for LSD, FMD and brucellosis with different transmission modes and rescaled *β* transmission coefficient values to isolate the effect of the mode of transmission. A, D, G (left) are DD models; B, E, H (centre) are FD models, and C, F, I (right) are DD models with rescaled *β* transmission of FD parameters. Rescaling LSD (A-B) and FMD (D-E) parameters have limited impact over the period modelled, but rescaling the brucellosis *β* shows a reduction in FD transmission.

**Figure 3.**
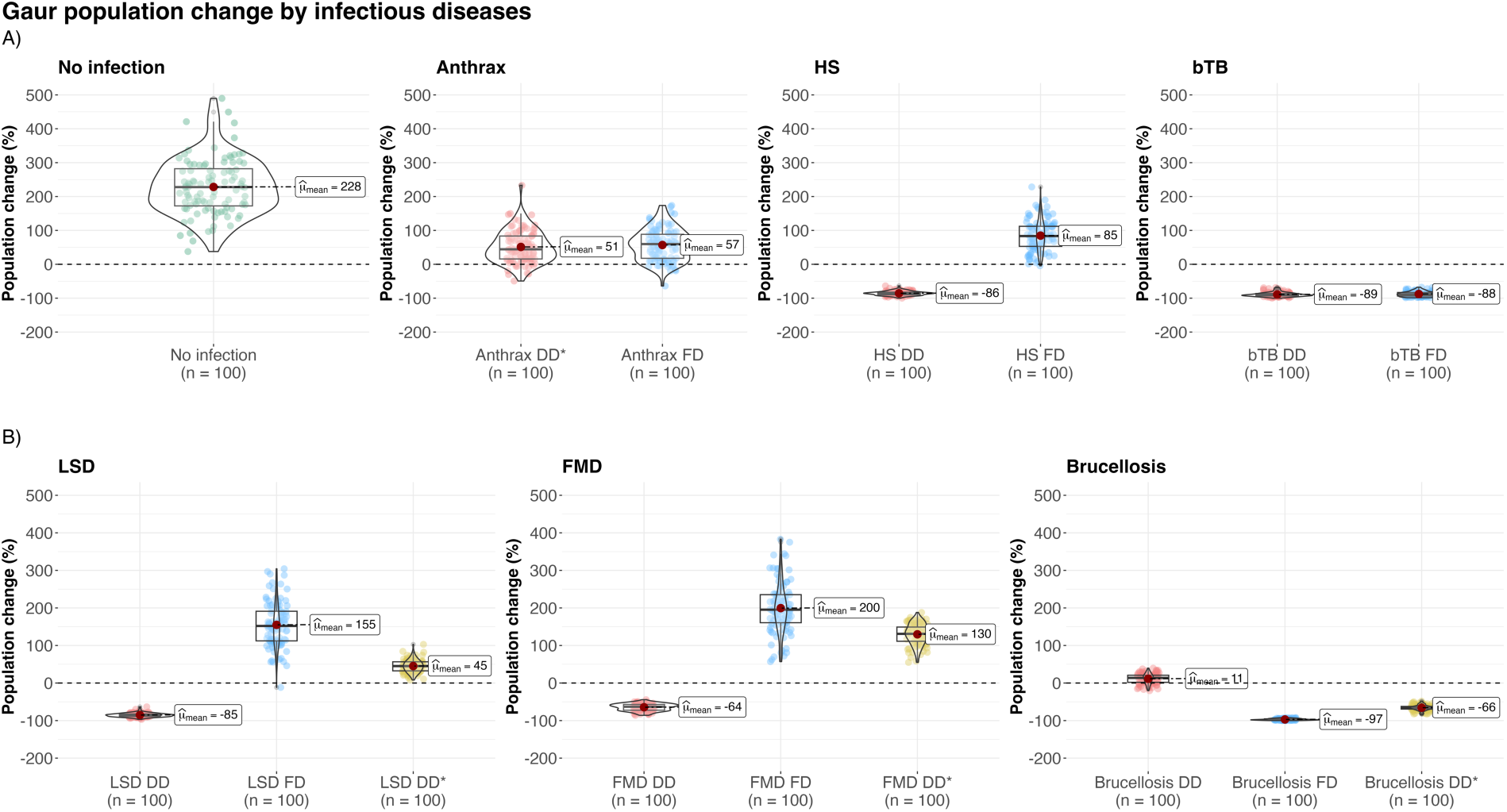
Overall modelled gaur population changes for each infection. Shown are the 100 results after 100 years of 100 stochastic simulations. The x-axis is the type of disease transmission and y axis is the population change in percentage. A) (top) compares no infection, FD and DD for anthrax, HS and bTB; B) (bottom) compares FD and DD transmission and DD* that uses the rescaled *β* transmission from the FD model with DD transmission for LSD, FMD, and brucellosis.

For LSD, FMD and brucellosis highlighting differences in population trends between FD and DD transmissions. We selected some important model results in Figure 1, and all modelling results and diagrams for infectious diseases and population changes can be found in the supplementary material, Figure S2 – Figure S16.

### Anthrax

There is no substantial impact on the population after introducing anthrax into the population, with a similar population change observed between FD and DD models. Both transmission rates showed an increase in population, with a 57% increase for FD and 51% for rescaled DD model. No massive die offs were predicted, with only 1-3 infectious animals predicted for each outbreak for both transmission modes, consistent with the low transmission rate (*β* = 0.01) applied and a rare case of animal-to-animal transmission.

### Bovine tuberculosis

There was uncertainty regarding the mode of transmission in bTB models, however, (31) showed limited qualitative differences in model outcomes when they used FD or DD transmission. Here, we saw similar results in that overall the populations tended to decline gradually through the simulation period with a 88-89% decline from the initial population. Rescaling the *β* transmission parameter also led to limited qualitative differences in population trends, but we did see differences in predicted classes; for example, this increased or decreased the number of infected individuals over time (i.e., higher or lower prevalence) (see supplementary materials, Figure S5 – S6).

### Haemorrhagic septicaemia

For HS, we found that the impact of infection was less dependent on the mode of transmission than case fatality. A ten-fold increase in fatality rate led to a decline in the total population change (Figure S8 – S9).

### Lumpy skin disease

For LSD, we found that the two published transmission parameter values (0.008 and 0.032) led to differing outcomes that also depended on the mode of transmission (41). Whilst rescaling the parameters did not lead to qualitative differences, the use of the parameter values estimated from direct density-dependent transmission within herds from (41) led to a population decline. However, the estimate from the indirect transmission (presumably via mechanical transmission from flies) did not and the modelled population still grew by 155% with FD LSD (Figure S11).

### Foot and mouth disease

The least impact on the modelled population was seen in the FMD model with FD transmission, which predicted the total population growing by 200%, around 28% less than the disease-free population. Frequency-dependent FMD transmission with a *β* transmission rate of 0.115 and the rescaled DD parameter 3e-4 similarly had limited impact on the population growth with an increasing population over time (Figure 2). Increasing *β* in the DD model, however, decreased the total population by-80% at *β* = 21, which had a greater impact on the population change from 130% at *β* = 3e-4. FMD also showed a periodic pattern with outbreaks around every 3-5 years (Figure 1). Increasing the *β* rate from 0.11 to 21 in FD FMD models led to similar dynamics close to DD transmission (Figure S13 – Figure S14).

### Brucellosis

Brucellosis with FD transmission led to a 97% decrease in the average population change (Figure 2 (H)) and was most likely to drive the population to local extinction with 16% of the total simulations leading to extinction, mostly occurring from year 80-100 (Figure 1 (H, P), and Figure S15).

## Discussion

Interactions between wildlife and livestock can facilitate the transmission of emerging infectious diseases (75), making this interface an essential area of concern to public health, animal production and wildlife conservation. We identified the potential consequences and severity of six bovine infectious diseases present in Thailand (anthrax, HS, bTB, LSD, FMD and brucellosis) in a model wild bovid population, using different infectious disease model compartments based on the current literature (Table 2). Brucellosis had the greatest population impacts and FMD the lowest, despite the same model structures being used for these two pathogens. Overall, our base model predicted population growth with varying impacts of diseases, and our analyses matched our expectation that those acute infections with very high fatality rates (anthrax and HS) have less impact than chronic infections with lower infectious rates (bTB, brucellosis), as infected individuals are rapidly removed from populations (10). Therefore, our analyses suggest that pathogens with longer incubation periods, chronic infection and low to medium fatality rates have a greater negative impact on population growth in small populations of endangered bovids (Figure 1 – Figure 3, Figure S5 – S6 and S15). This is most likely because these traits allow infections to persist, allowing long-term infection effects on demographic structures (e.g. reduced birth rate, increased death rate).

We used 100% fatality rates in all infected animals as the worst-case scenario for the anthrax model, which led to limited impact over the 100 years, likely because of this rapid removal of infected individuals (*I*), despite repeated reintroduction of infection (10). We first considered anthrax transmission between infected and susceptible animals as FD transmission, assuming contact rate is more influential than host density (29). However, the transmission mode could also be DD, based on the density of spores in the contaminated environment (e.g. infected carcass, soil) (28) and the cattle density that could contribute to between-species (76). Thus, we modelled repeated introductions through *ɛ* to cover the external force of infections, including the risk of disease transmission from cattle, other than within-herd transmission.

Bovine tuberculosis causes chronic, fatal infection and reduces pregnancy rates and, therefore, the population growth of wild bovids (32, 77). There is no current evidence of bTB infection driven population declines in Asian wild bovids, however our study found that, regardless of both transmission modes, the long-term effect of bTB would be to reduce the expected total population by around 88-89%. This is similar to findings by Jolles et al. (2005) (32), who showed that bTB persisted in African buffalo populations and reduced adult buffalo numbers primarily through mortality of animals more than 4.5 years old. The transmission coefficient (*β*) was noted as one of the most important parameters for bTB in African buffalo (31). In our work, we found consistent population dynamics between FD and DD transmission, defined by similar trends and percentages of population change, when converting *β* between the original value (from several studies) and the rescaled values (Figure 2 – Figure 3). This is likely due to the duration of the infection, which might increase the probability of contact with infectious animals in the population and the number of transmissions.

For HS, many animals are infected but do not develop clinical signs, making it difficult to detect an infected animals. Further, variable clinical signs make positive cases difficult to detect, therefore the case fatality rate is normally substantially higher than the actual infection fatality rate. In our study, we calculated the fatality rate using the prevalence of seropositive animals (max = 40%) from reported studies of cattle populations, and this substantially decreased the case fatality from 90% to around 6% of animals (40, 78). Our model shows that changing the mortality from 0.53% to 5.8% affects the total population numbers more than changing the transmission modes, by more strongly reducing the population sizes (Figure S8 – Figure S9). HS antibodies were found in free-ranging buffalo in Asia so this population might be a reservoir, but this needs further investigation (37). HS is endemic among cattle in Thailand (78), and mortality in wild ungulates has been reported historically (79), so the mortality and infection status of HS should be considered in the mitigation plans for endangered species (e.g. wild water buffalo and banteng).

Both FMD and LSD with FD transmission had the least impact on populations, with acute, short infections with lower overall mortality (74, 80). FMDV in particular, is highly contagious among cattle with a very high *β* coefficient compared to the other diseases (27). Yet, although Beck-Johnson, Gorsich (27) found little effect of FMD with either transmission modes within-herds, our results showed that DD transmission led to greater population declines, as did rescaling the parameter used for FD models. The reason for the latter observation is not clear but might be because the dynamics with reintroduction allow more infection to persist and so suppress the population (Figure 2). Note that with reported wild bovid herd sizes, acute transmission is unlikely to allow FMDV to persist, but reintroductions from cattle reservoirs (modelled through *ɛ*) are likely (73). Our result for FMD DD transmission also showed cyclic patterns in outbreaks consistent with seasonal patterns of outbreaks observed in Thailand (81).

We found that brucellosis with FD transmission and its reported *β* rate might cause extinction 16% of the time, whereas DD transmission (*β* = 5e-6) may suppress population growth, but not enough to cause population declines and even with the published FD rate rescaled (*β* = 1e-5) and used in a DD model, this caused declines but not extinctions (Figure 3; (B)). Brucellosis has caused population declines among African buffalo, especially when there is co-infection with tuberculosis (82). However, brucellosis only caused limited population growth impacts in American bison, even though the disease persisted in the population over time (61, 83). In Dobson and Meagher’s study (61), their FD brucellosis models showed bison populations would increase in numbers, whereas our models predicted a decrease, perhaps because our model species’ population size and structure differed from their study. Notably, *Brucella* can infect multiple species, and the transmission source may not be obvious when multiple species interact. For example, brucellosis outbreaks in Yellowstone National Park, USA, were not from wild bison as first thought, with elk the likely primary host (84). Understanding the potential transmission among and from other wild Asian ungulates may be necessary to fully understand potential brucellosis impacts.

We assumed a single, closed (no migration) population with constant natural birth and death rates. Therefore, our models explore the intrinsic population dynamics without considering the influence of other positive (e.g. conservation) or negative factors (e.g. habitat destruction, competition). Furthermore, it is unclear what population changes occur during migration (85), so a closed population model can only simulate within-herd dynamics and reflect the population impacts in a small population, such as in small protected areas (20, 22).

Selecting the appropriate transmission mode for modelling is challenging (86). The infectious disease parameter values themselves are mostly estimated from livestock outbreak data, which can vary among the regions. Rarely is infection ‘natural’ without intervention through disease control (27). Although the transmission type for some pathogens have been recorded as FD or DD in previous studies, these were mainly conducted under farm husbandry or experimental conditions in captive or closed systems. These conditions significantly allow animal density to affect contact rates. However, our study focused on wildlife populations that are distributed in areas in which the frequency of contact could have more influence. Moreover, some infectious diseases can display aspects of FD and DD depending on the conditions, such as within-or between herd transmission, herd size, density, contact with other reservoirs and contact mode (indirect, direct). For example, the bTB transmission rate can be increased correlated to herd size if the area is stable because the density of animals is increased (87). Also, the transmission mode for anthrax spores from animal to animal is FD, but from the environment to an animal is based on the density of the spores in the areas. We, therefore, took the strategy of assuming the most extreme scenarios, fully FD and DD, and used both for modelling.

Our models also added the external force of infection (*ɛ*), which represents the re-introduction of pathogens. *ɛ* is assumed to include transmission from other sources of infection other than just infectious animals, such as transmission due to environmental factors (e.g. soil, carcasses) or vectors (e.g. blood-sucking fly) to susceptible animals (10). This transmission can theoretically cause population extinctions if agents have high case fatality rates. Here, we chose a relatively high reintroduction rate (∼ 2 per year into the initial population), which likely represents a worst-case scenario. However, to improve this study, we encourage adding the specific environmental factors for each disease and incorporating spatial analyses (88, 89).

Further studies might also consider adding the potential reservoir hosts and their dynamics into the models by building two or more host models to examine the transmission route among the potential hosts (90–94). Modelling coinfection is another important point as there are interactions between infections such as FMDV and HS, which seen as a secondary infection in FMD outbreaks (95), or between brucellosis and bTB (82). However, our analyses provide an approach to understanding the *relative* likely impact of common endemic and emerging diseases with different traits and is a tool for understanding gaps in disease surveillance and control systems by using the prediction modelling before implementing actions. Future analyses could also determine the impact of using an Erlang-distributed waiting time, rather than an exponential distribution, on those parameters with large amounts of variation, particularly the incubation period (96). Another further analysis is a sensitivity analysis that can be applied to identify the degree of influence of the disease parameters on the model output, in this case, population change. It also suggests which state of disease transmission should prompt action and aids in selecting optimal control measures (97).

Strengthening disease surveillance and mitigation programs may be further achieved by targeting virulent diseases through passive and active surveillance data, such as collecting the frequency of infections, number and species of wild ungulates, behaviour and time spent together between wild and domestic livestock (particularly in the high-risk areas) (98, 99). It may be useful for disease mitigation to largely focus on domestic animal disease control and preventing transmission to wildlife as an amenable approach (98). Moreover, conserving wildlife habitat can reduce the probability of contact and the risk of disease transmission between wildlife and domestic livestock (3, 100). Limiting the contact between wildlife and livestock could reduce species extinction (101).

With applications in wildlife conservation, a reproducible modelling framework is advantageous for targeting pathogens that threaten other wildlife populations with similar assumptions. Although our infectious disease modelling focused on the traits of pathogens in one species population, our method and framework may be applicable to other wildlife populations by incorporating their population demographics and disease parameters. This framework is also beneficial for endangered species, enabling the simulation of various scenarios and the identification of potential disease threats, along with estimating the recovery period after introducing the infection.

## Conclusion

Our study has provided a prediction of the potential consequence of disease in wild bovid populations considering six important bovine infectious diseases; anthrax, HS, bTB, LSD, FMD and brucellosis. The baseline population model shows a natural population growth of ∼ 228%, suggesting maintaining healthy vulnerable populations could allow them to reestablish and overcome current levels of extinction threats while diseases and other factors may regulate population growth. The inclusion of different disease traits has consequences on the population numbers depending on the transmission, incubation, fatality and infectious rates. Brucellosis and bTB models show the greatest, long-term impact among all the models, whereas FMD and LSD showed the least impact, suggesting common but more chronic or’slow’ infections with relatively high mortality may pose the greatest threat to smaller, threatened bovid populations.

## Acknowledgements

This work was supported by funds from the New Zealand Ministry of Foreign Affairs and Trade Manaaki New Zealand Scholarship (WH), Royal Society Te Apārangi Rutherford Discovery Fellowship (RDF-MAU1701) and the Percival Carmine Chair in Epidemiology and Public Health (both DTSH). We thank Prof Richard Laven from Massey University for helpful comments on HS and Dr Rosemary Barraclough from *^m^*EpiLab for revising the manuscript draft. The authors thank Massey University’s subscription to New Zealand eScience Infrastructure (NeSI) which enabled us to use high-performance computing facilities https://www.nesi.org.nz.

