## Supplementary materials for "Impact of Infectious Diseases on Wild Bovidae Populations in Thailand: Insights from Population Modelling and Disease Dynamics"

Includes:

- Population dynamics model results (Figure S1)
- Infectious disease model structures and results (Figure S2 – S16)
- PCA biplot (Figure S17-S18)

**Abbreviation**

| Symbol | Description |
|---|---|
| $N_0$ | Starting total population |
| $S$ | Susceptible |
| $E$ | Exposed |
| $I$ | Infected |
| $R$ | Recovered |
| $M$ | Calves with maternally derived immunity |
| $a$ | Adult |
| $sa$ | Subadult |
| $C$ | Calf |
| $\mu_b$ | Birth rate |
| $\mu_a$ | Adult death rate |
| $\mu_{sa}$ | Subadult death rate |
| $\mu_c$ | Calf death rate |
| $\delta$ | Ageing rate |
| $\beta$ | Disease transmission rate |
| $\sigma$ | 1/Incubation period |
| $\gamma$ | Infectious period |
| $\rho$ | Disease-induced fatality |
| $\alpha$ | Infected female will produce infected calf |
| $\mu_{bI}$ | Birth rate for infectious individuals |
| $\omega$ | Loss of immunity for calves |
| $\omega_m$ | Waning of maternal immunity |
| $\epsilon$ | External force of infection rate |
| * | Rescaling of the transmission mode |

### Model structures and results

All models used three age structures (adult, subadult and calf). For more details of the disease parameters and values, see the main text.

### 1. Population dynamics

The population dynamics model included demography (birth, death and ageing rate), and no infection importation.

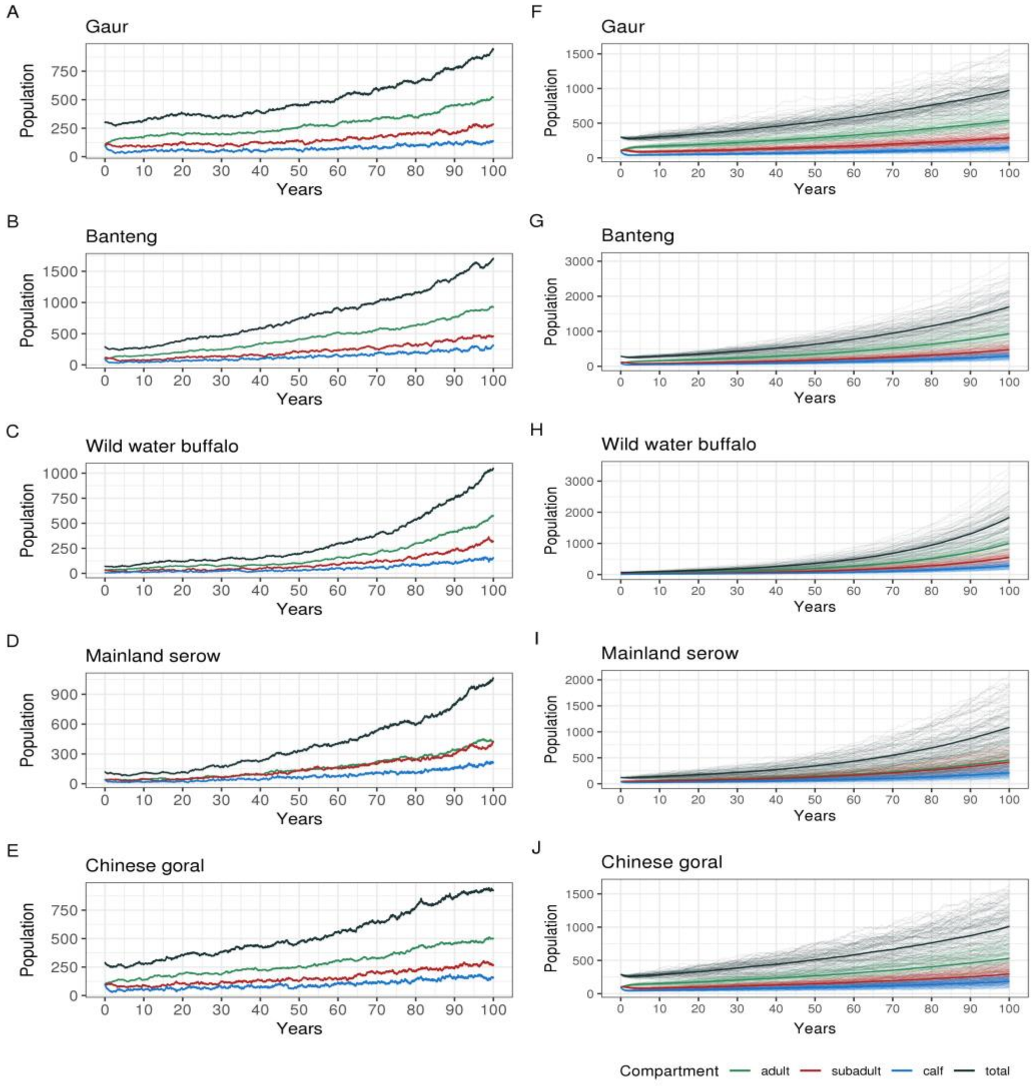


Figure S1 Single and 100 stochastic simulations of population dynamics for five wild bovids. A-E are single simulations; F-J are 100 simulations.

### 2. Infectious disease models

The population dynamics model with demography and infection importation (using infectious ($I$) = 1 for all models). The rescaled (*) $\beta$ rates were used to examine the consistency of the results between frequency-dependent (FD) and density-dependent (DD).

#### Anthrax (*Bacillus anthracis*)

The $SI$ model for anthrax with a 100% fatality rate (all infected will die).

$$
\begin{aligned}
N &= S_c + I_c + S_{sa} + I_{sa} + S_a + I_a \\
\frac{dS_c}{dt} &= \mu_b S_a - \beta_c S_c (I_c + I_{sa} + I_a) - \delta_c S_c - \mu_c S_c - \epsilon S_c \\
\frac{dS_{sa}}{dt} &= -\beta_{sa} S_{sa} (I_c + I_{sa} + I_a) + \delta_c S_c - \delta_{sa} S_{sa} - \mu_{sa} S_{sa} - \epsilon S_{sa} \\
\frac{dS_a}{dt} &= -\beta_a S_a (I_c + I_{sa} + I_a) + \delta_{sa} S_{sa} - \mu_a S_a - \epsilon S_a \\
\frac{dI_c}{dt} &= \beta_c S_c (I_c + I_{sa} + I_a) - \rho_c I_c + \epsilon S_c \\
\frac{dI_{sa}}{dt} &= \beta_{sa} S_{sa} (I_c + I_{sa} + I_a) - \rho_{sa} I_{sa} + \epsilon S_{sa} \\
\frac{dI_a}{dt} &= \beta_a S_a (I_c + I_{sa} + I_a) - \rho_a I_a + \epsilon S_a
\end{aligned} \tag{1}
$$

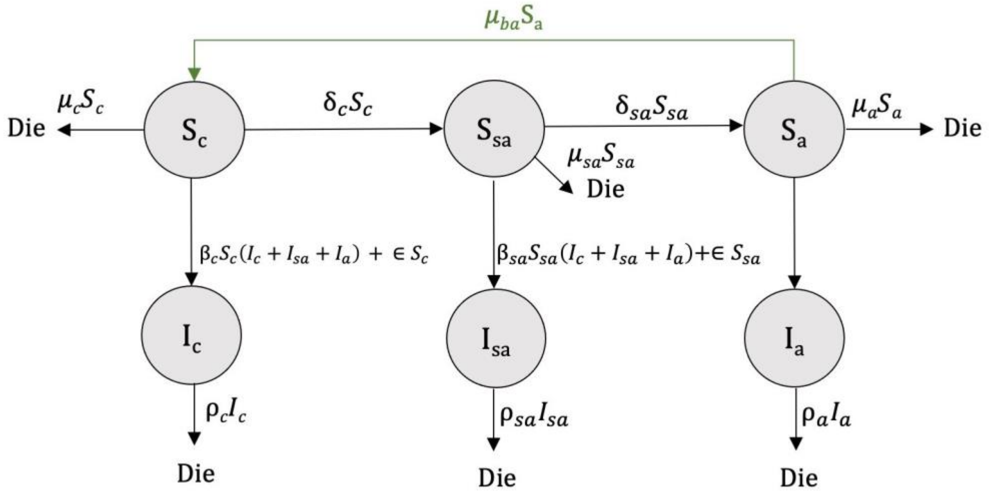


Figure S2 $SI$ model diagram of anthrax.

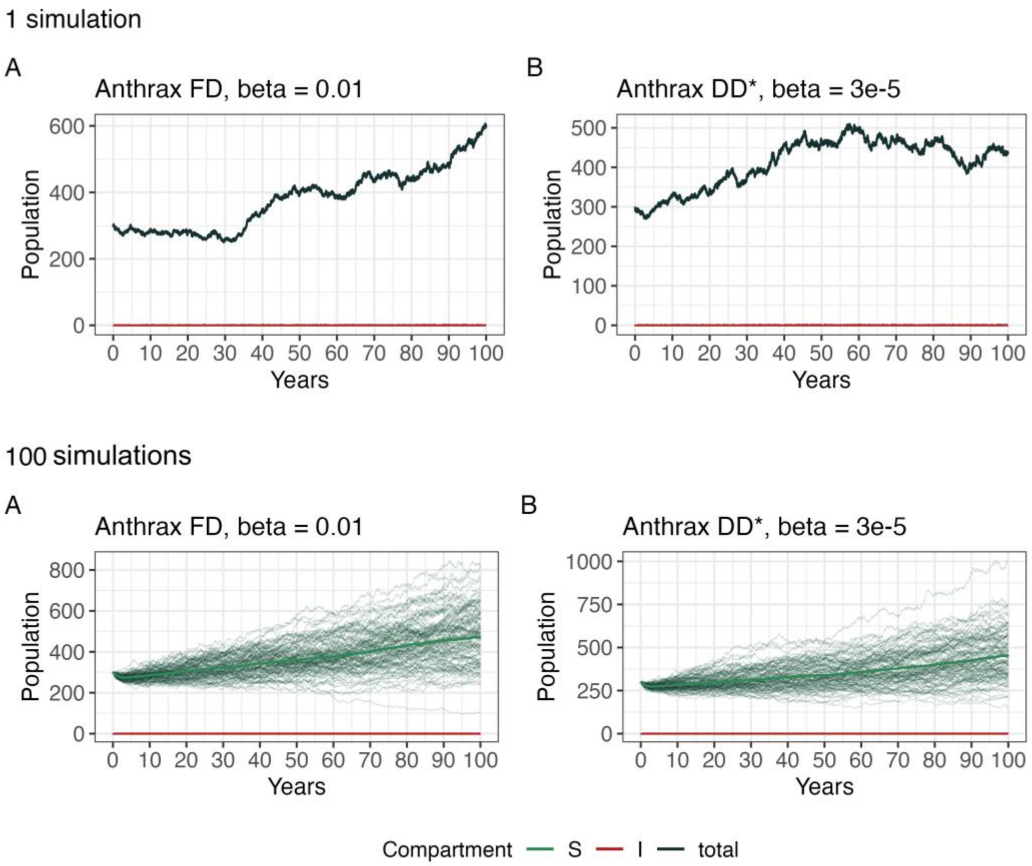


Figure S3 Single and 100 stochastic simulations of anthrax show that the population has increased with a few outbreak events, indicating rare animal-to-animal transmission. Changing the $\beta$ rate (from 0.01 to 3e-5) slightly affects the total population.

**Bovine tuberculosis (bTB - *Mycobacterium tuberculosis*)**

The $SEI$ model for bTB included a lifelong infection with no recovery state. Therefore, an infected individual will remain in the infectious state until die either from disease-induced or natural death.

$$
\begin{aligned}
N &= S_c + E_c + I_c + S_{sa} + E_{sa} + I_{sa} + S_a + E_a + I_a \\
\frac{dS_c}{dt} &= \mu_b(S_a + E_a) + \mu_{bI} I_a - \beta_c S_c(I_c + I_{sa} + I_a) - \delta_c S_c - \mu_c S_c - \epsilon S_c \\
\frac{dS_{sa}}{dt} &= -\beta_{sa} S_{sa}(I_c + I_{sa} + I_a) + \delta_c S_c - \delta_{sa} S_{sa} - \mu_{sa} S_{sa} - \epsilon S_{sa} \\
\frac{dS_a}{dt} &= -\beta_a S_a(I_c + I_{sa} + I_a) + \delta_{sa} S_{sa} - \mu_a S_a - \epsilon S_a \\
\frac{dE_c}{dt} &= \beta_c S_c(I_c + I_{sa} + I_a) - \sigma_c E_c - \delta_c E_c - \mu_c E_c + \epsilon S_c \\
\frac{dE_{sa}}{dt} &= \beta_{sa} S_{sa}(I_c + I_{sa} + I_a) - \sigma_{sa} E_{sa} + \delta_c E_c - \delta_{sa} E_{sa} - \mu_{sa} E_{sa} + \epsilon S_{sa} \\
\frac{dE_a}{dt} &= \beta_a S_a(I_c + I_{sa} + I_a) - \sigma_a E_a + \delta_{sa} E_{sa} - \mu_a E_a + \epsilon S_a \\
\frac{dI_c}{dt} &= \sigma_c E_c - (\rho_c + \mu_I) I_c - \delta_c Ic \\
\frac{dI_{sa}}{dt} &= \sigma_{sa} E_{sa} - (\rho_{sa} + \mu_{sa}) I_{sa} + \delta_c Ic - \delta_{sa} Isa \\
\frac{dI_a}{dt} &= \sigma_a E_a - (\rho_{sa} + \mu_a) I_a + \delta_{sa} Isa
\end{aligned} \tag{2}
$$

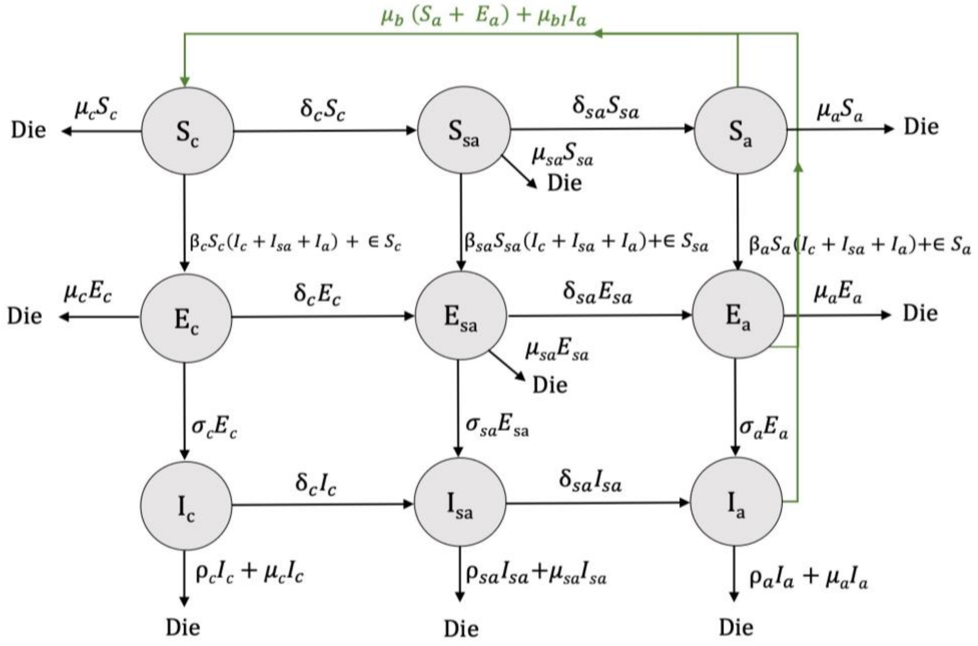


Figure S4 $SEI$ model diagram of bTB.

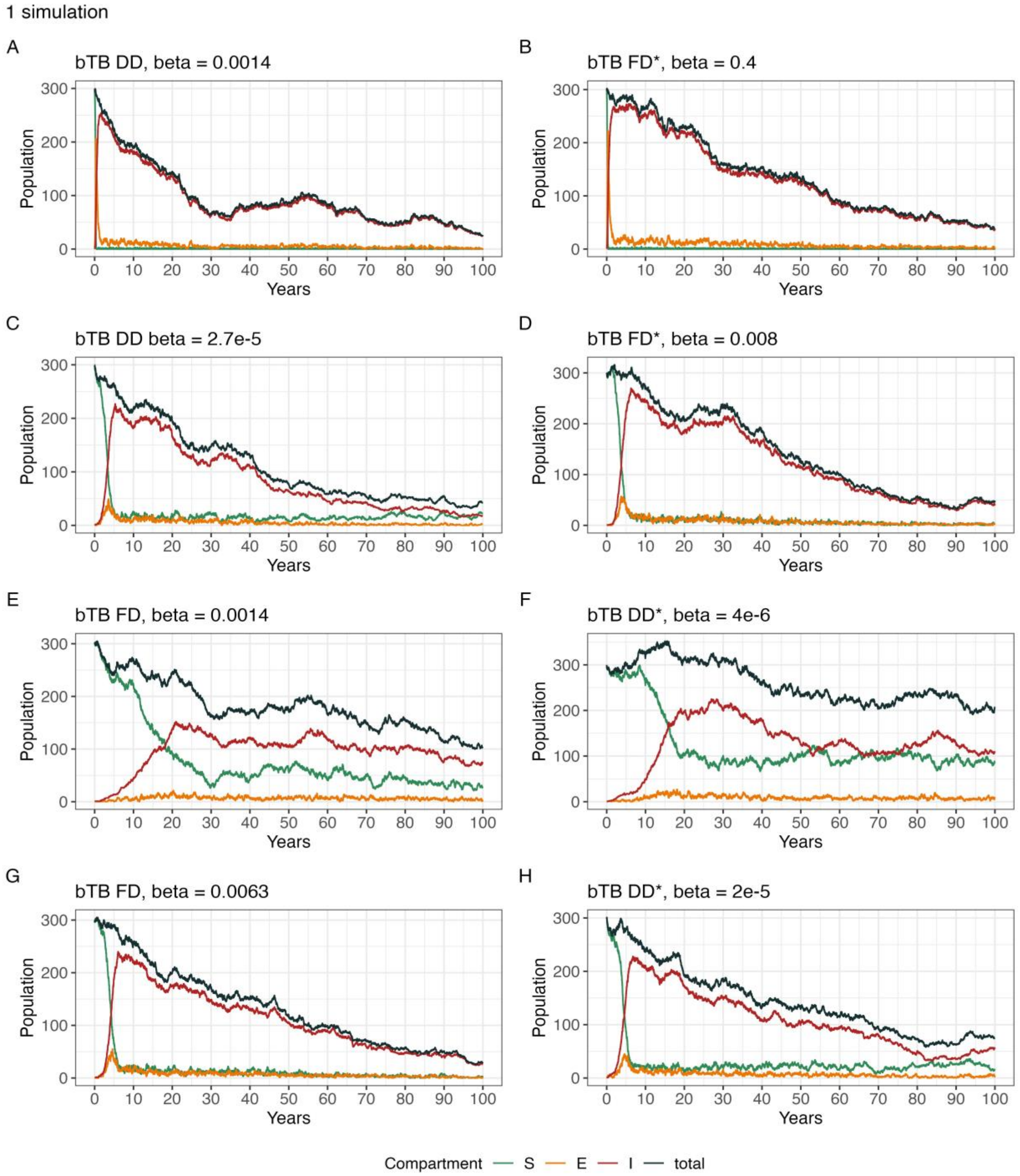


Figure S5 Single stochastic simulations of bTB. During the initial period after infection, susceptible animals decreased as they became exposed and then infected. Then, the population decreased over time due to chronic infection without recovery.

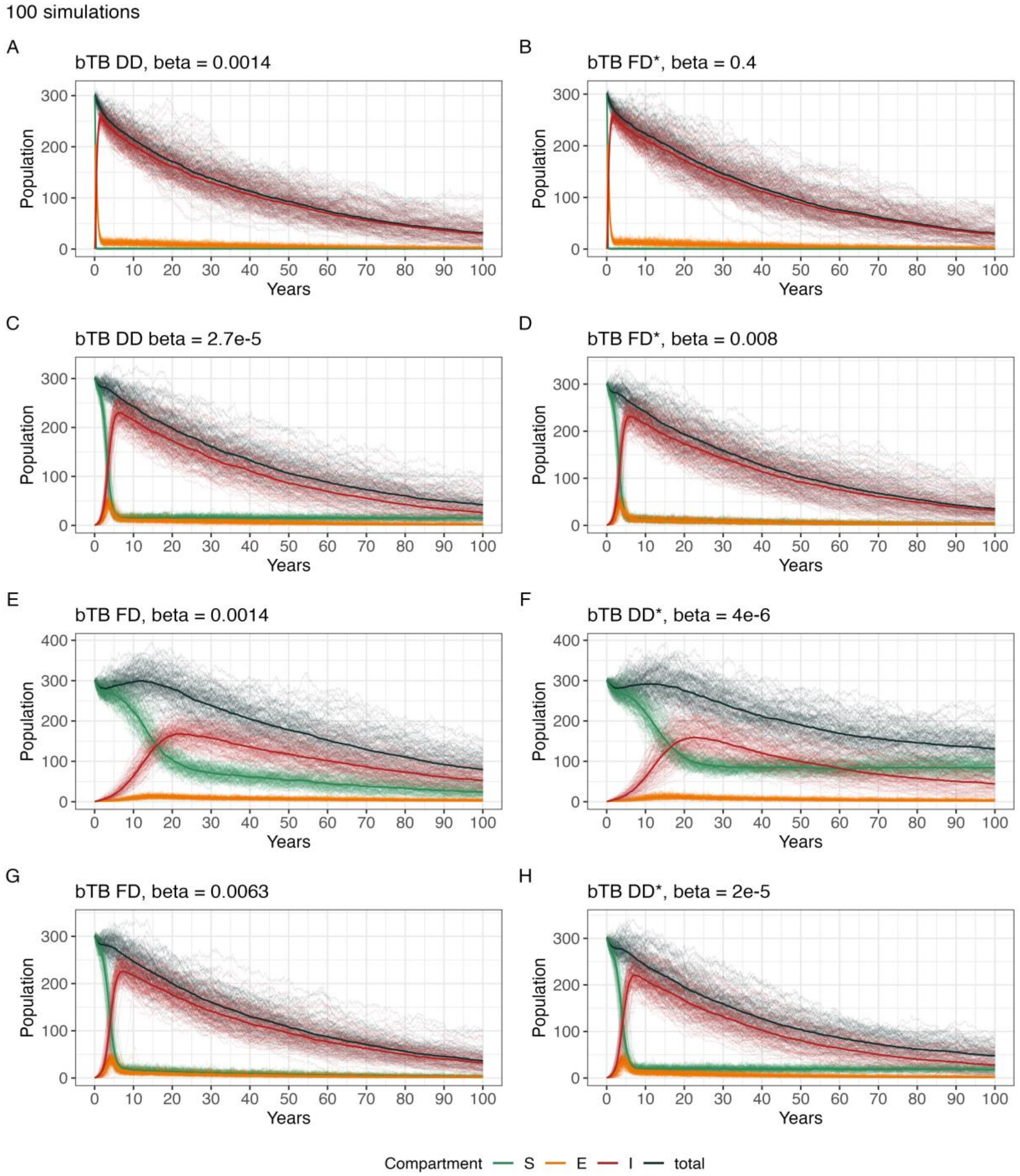


Figure S6 Hundred stochastic simulations of bTB.

**Hemorrhagic septicaemia (HS - *Pasteurella multocida*)**

The $SIRS$ model for HS used two fatality rates, 0.5% and 5%.

$$
\begin{aligned}
N &= S_c + I_c + R_c + S_{sa} + I_{sa} + R_{sa} + S_a + I_a + R_a \\
\frac{dS_c}{dt} &= \mu_b N_a - \beta_c S_c (I_c + I_{sa} + I_a) - \delta_c S_c - \mu_c S_c + \omega_c R_c - \epsilon S_c \\
\frac{dS_{sa}}{dt} &= -\beta_{sa} S_{sa} (I_c + I_{sa} + I_a) + \delta_c S_c - \delta_{sa} S_{sa} - \mu_{sa} S_{sa} + \omega_{sa} R_{sa} - \epsilon S_{sa} \\
\frac{dS_a}{dt} &= -\beta_a S_a (I_c + I_{sa} + I_a) + \delta_{sa} S_{sa} - \mu_a S_a + \omega_a R_a - \epsilon S_a \\
\frac{dI_c}{dt} &= \beta_c S_c (I_c + I_{sa} + I_a) - (1 - \rho_c)\gamma_c I_c - \rho_c \gamma_c I_c - \delta_c I_c - \mu_c I_c + \epsilon S_c \\
\frac{dI_{sa}}{dt} &= \beta_{sa} S_{sa} (I_c + I_{sa} + I_a) - (1 - \rho_{sa})\gamma_{sa} I_{sa} - \rho_{sa}\gamma_{sa} I_{sa} + \delta_c I_c - \delta_{sa} S_{sa} - \mu_{sa} I_{sa} + \epsilon S_{sa} \\
\frac{dI_a}{dt} &= \beta_a S_a (I_c + I_{sa} + I_a) - (1 - \rho_a)\gamma_a I_a - \rho_a \gamma_a I_a + \delta_{sa} S_{sa} - \mu_a I_a + \epsilon S_a \\
\frac{dR_c}{dt} &= (1 - \rho_c)\gamma_c I_c - \omega_c R_c - \delta_c R_c - \mu_c R_c \\
\frac{dR_{sa}}{dt} &= (1 - \rho_{sa})\gamma_{sa} I_{sa} - \omega_{sa} R_{sa} + \delta_c R_c - \delta_{sa} R_{sa} - \mu_{sa} R_{sa} \\
\frac{dR_a}{dt} &= (1 - \rho_a)\gamma_a I_a - \omega_a R_a + \delta_{sa} R_{sa} - \mu_a R_a
\end{aligned} \tag{3}
$$

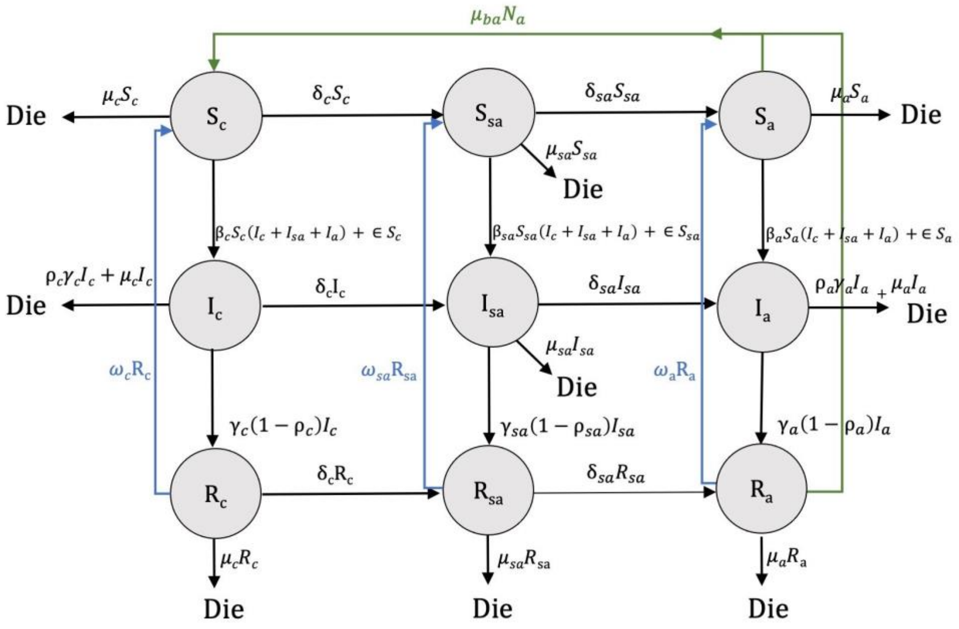


Figure S7 $SIRS$ model diagram of HS.

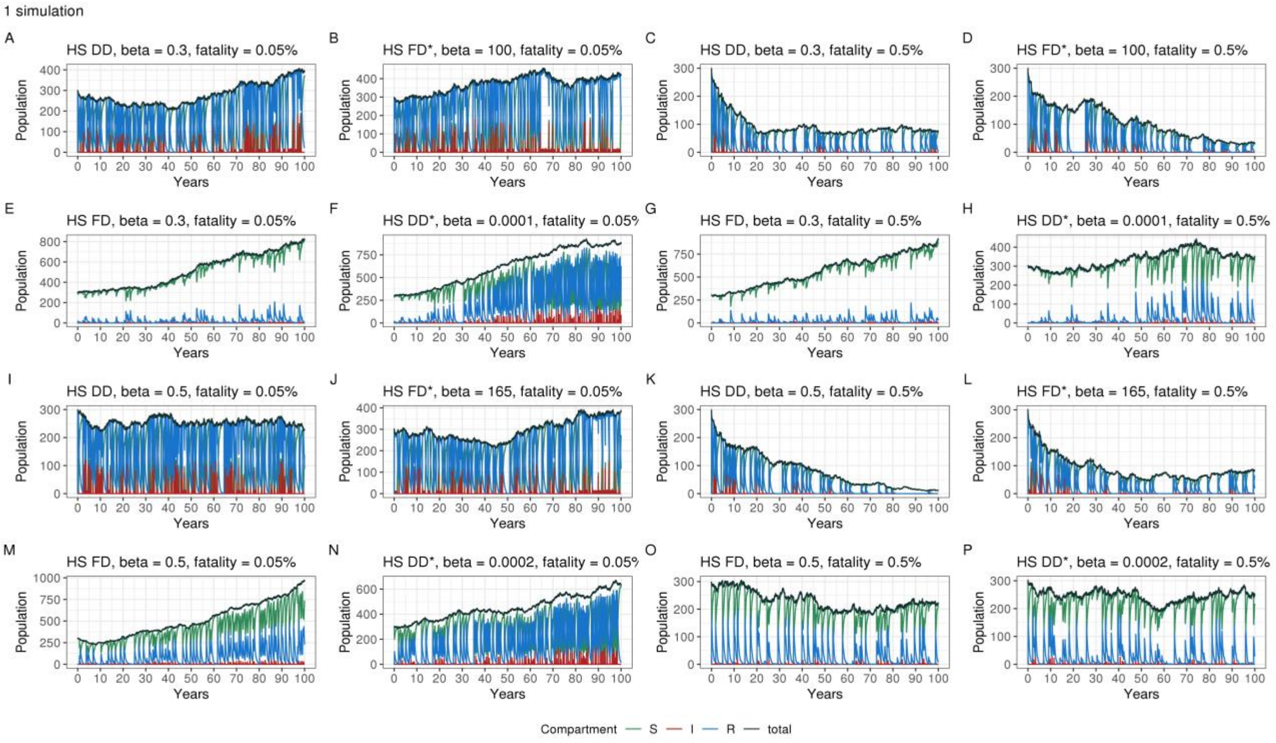


Figure S8 Single stochastic simulations of HS show that increasing the $\beta$ rate (0.3 → 0.5) affects the population number for DD models (A, I) but not for FD models (E, M). However, increasing the fatality rate (0.05% → 0.5%) shows a significant impact on DD models for both $\beta$ rates (A, C & I, K) and the FD model for a 0.5 transmission rate (K).

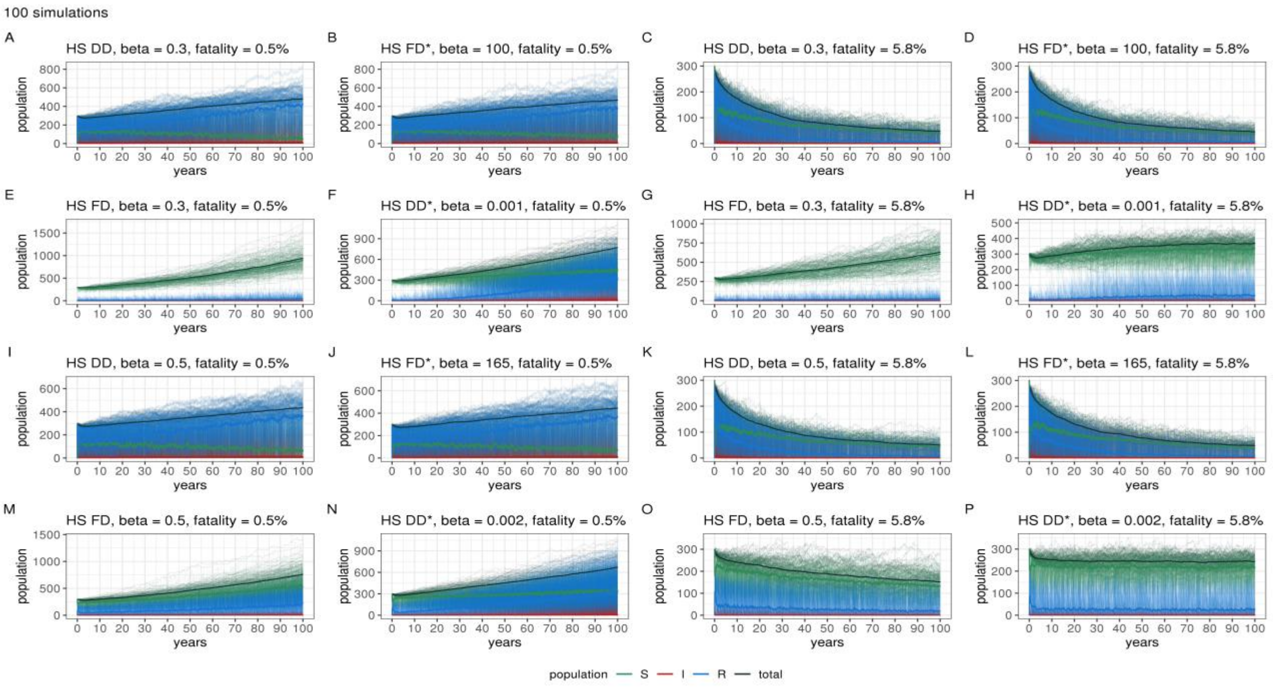


Figure S9 Hundred stochastic simulations of HS.

**Lumpy skin disease (LSD - Capripoxvirus)**

The *SEIRS* model for LSD included the exposed and recovery state with the birth rate for infected mothers.

$$
\begin{aligned}
N &= S_c + E_c + I_c + R_c + S_{sa} + +E_{sa} + I_{sa} + R_{sa} + S_a + E_a + I_a + R_a \\
\frac{dS_c}{dt} &= \mu_b(S_a + E_a + R_a) + \mu_b I_a - \beta_c S_c(I_c + I_{sa} + I_a) - \delta_c S_c - \mu_c S_c + \omega_c R_c - \epsilon S_c \\
\frac{dS_{sa}}{dt} &= -\beta_{sa} S_{sa}(I_c + I_{sa} + I_a) + \delta_c S_c - \delta_{sa} S_{sa} - \mu_{sa} S_{sa} + \omega_{sa} R_{sa} - \epsilon S_{sa} \\
\frac{dS_a}{dt} &= -\beta_a S_a(I_c + I_{sa} + I_a) + \delta_{sa} S_{sa} - \mu_a S_a + \omega_a R_a - \epsilon S_a \\
\frac{dE_c}{dt} &= \beta_c S_c(I_c + I_{sa} + I_a) - \sigma_c E_c - \delta_c E_c - \mu_c E_c + \epsilon S_c \\
\frac{dE_{sa}}{dt} &= \beta_{sa} S_{sa}(I_c + I_{sa} + I_a) - \sigma_{sa} E_{sa} + \delta_c E_c - \delta_{sa} E_{sa} - \mu_{sa} E_{sa} + \epsilon S_{sa} \\
\frac{dE_a}{dt} &= \beta_a S_a(I_c + I_{sa} + I_a) - \sigma_a E_a + \delta_{sa} E_{sa} - \mu_a E_a + \epsilon S_a \\
\frac{dI_c}{dt} &= \sigma_c E_c - (1 - \rho_c)\gamma_c I_c - \rho_c \gamma_c I_c - \delta_c I_c - \mu_c I_c \\
\frac{dI_{sa}}{dt} &= \sigma_{sa} E_{sa} - (1 - \rho_{sa})\gamma_{sa} I_{sa} - \rho_{sa}\gamma_{sa} I_{sa} + \delta_c I_c - \delta_{sa} I_{sa} - \mu_{sa} I_{sa} \\
\frac{dI_a}{dt} &= \sigma_a E_a - (1 - \rho_a)\gamma_a I_a - \rho_a \gamma_a I_a + \delta_{sa} S_{sa} - \mu_a I_a \\
\frac{dR_c}{dt} &= (1 - \rho_c)\gamma_c I_c - \omega_c R_c - \delta_c R_c - \mu_c R_c \\
\frac{dR_{sa}}{dt} &= (1 - \rho_{sa})\gamma_{sa} I_{sa} - \omega_{sa} R_{sa} + \delta_c R_c - \delta_{sa} R_{sa} - \mu_{sa} R_{sa} \\
\frac{dR_a}{dt} &= (1 - \rho_a)\gamma_a I_a - \omega_a R_a + \delta_{sa} R_{sa} - \mu_a R_a
\end{aligned} \tag{4}
$$

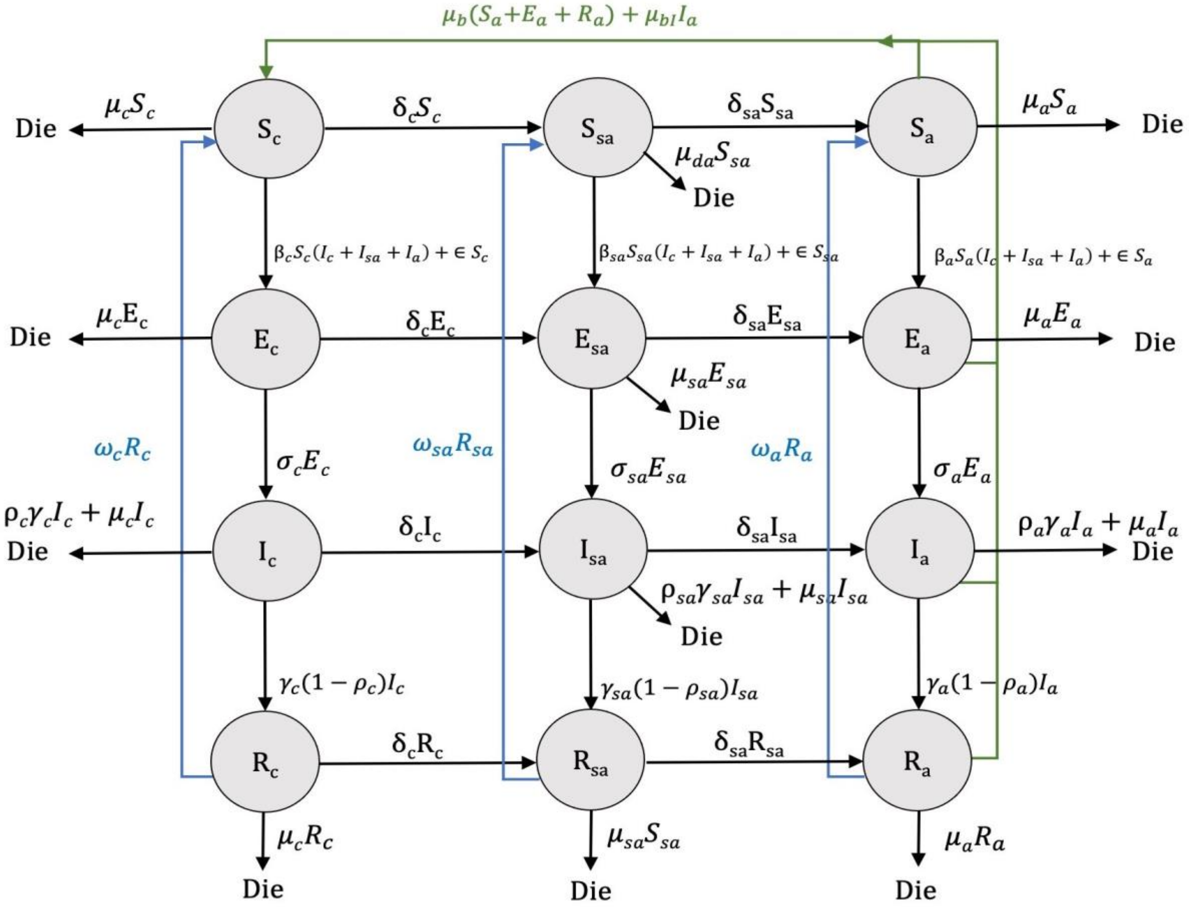

Figure S10 $SEIRS$ model diagram of LSD.

1 simulation

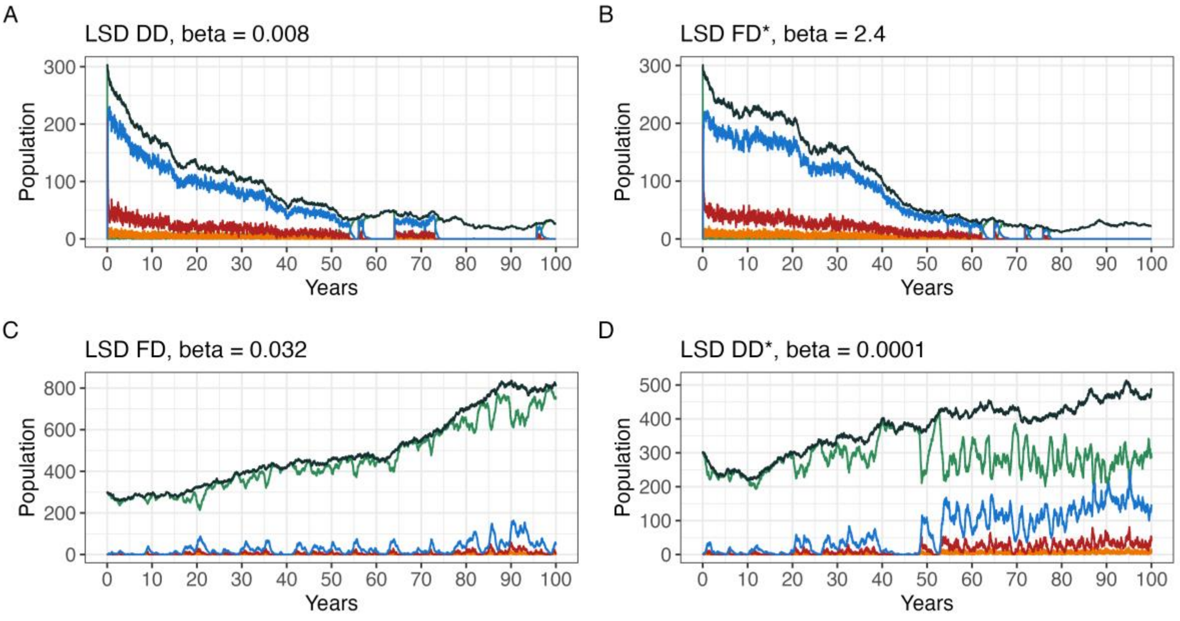


100 simulations

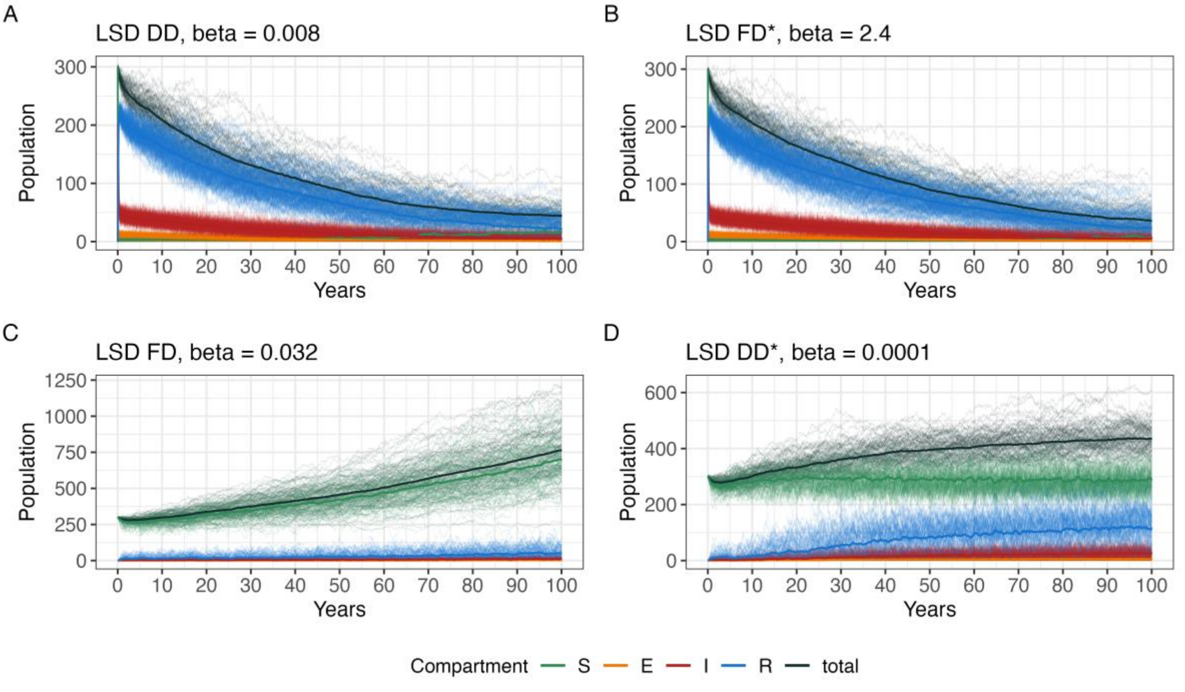


Figure S11 Single and 100 stochastic simulations of LSD. FD transmission had less impact on the populations than DD transmission, which resulted in a total population decrease over time.

**Foot and mouth disease (FMD - Aphthovirus) & Bovine Brucellosis (*Brucella abortus*)**

The *SEIRM/E* model for FMD and Brucellosis included calves with derived maternal immunity, susceptible calves after waning maternal immunity, and the effect on the birth rate of an infectious mother.

$$
\begin{aligned}
N &= S_c + E_c + I_c + R_c + S_{sa} + E_{sa} + I_{sa} + R_{sa} + S_a + E_a + I_a + R_a + M + S_m \\
\frac{dS_c}{dt} &= \mu_b(S_a + E_a) + (1-\alpha)\mu_b I_a - \beta_c S_c(I_c + I_{sa} + I_a) - \delta_c S_c - \mu_c S_c + \omega_c R_c - \epsilon S_c \\
\frac{dS_{sa}}{dt} &= -\beta_{sa} S_{sa}(I_c + I_{sa} + I_a) + \delta_c S_c - \delta_{sa} S_{sa} - \mu_{sa} S_{sa} + \omega_{sa} R_{sa} + \delta_m S_m - \epsilon S_{sa} \\
\frac{dS_a}{dt} &= -\beta_a S_a(I_c + I_{sa} + I_a) + \delta_{sa} S_{sa} - \mu_a S_a + \omega_a R_a - \epsilon S_a \\
\frac{dE_c}{dt} &= \beta_c S_c(I_c + I_{sa} + I_a) + \beta_c S_m(I_c + I_{sa} + I_a) - \sigma_c E_c - \delta_c E_c - \mu_c E_c + \epsilon S_c \\
\frac{dE_{sa}}{dt} &= \beta_{sa} S_{sa}(I_c + I_{sa} + I_a) - \sigma_{sa} E_{sa} + \delta_c E_c - \delta_{sa} E_{sa} - \mu_{sa} E_{sa} + \epsilon S_{sa} \\
\frac{dE_a}{dt} &= \beta_a S_a(I_c + I_{sa} + I_a) - \sigma_a E_a + \delta_{sa} E_{sa} - \mu_a E_a + \epsilon S_a \\
\frac{dI_c}{dt} &= \sigma_c E_c - (1-\rho_c)\gamma_c I_c - \rho_c \gamma_c I_c - \delta_c I_c - \mu_c I_c + \alpha \mu_{bI} I_a \\
\frac{dI_{sa}}{dt} &= \sigma_{sa} E_{sa} - (1-\rho_{sa})\gamma_{sa} I_{sa} - \rho_{sa}\gamma_{sa} I_{sa} + \delta_c I_c - \delta_{sa} I_{sa} - \mu_{sa} I_{sa} \\
\frac{dI_a}{dt} &= \sigma_a E_a - (1-\rho_a)\gamma_a I_a - \rho_a \gamma_a I_a + \delta_{sa} S_{sa} - \mu_a I_a \\
\frac{dR_c}{dt} &= (1-\rho_c)\gamma_c I_c - \omega_c R_c - \delta_c R_c - \mu_c R_c \\
\frac{dR_{sa}}{dt} &= (1-\rho_{sa})\gamma_{sa} I_{sa} - \omega_{sa} R_{sa} + \delta_c R_c - \delta_{sa} R_{sa} - \mu_{sa} R_{sa} \\
\frac{dR_a}{dt} &= (1-\rho_a)\gamma_a I_a - \omega_a R_a + \delta_{sa} R_{sa} - \mu_a R_a \\
\frac{dM}{dt} &= \mu_b R_a - \omega_M M - \mu_c M \\
\frac{dS_m}{dt} &= \omega_M M - \delta_m S_m - \beta_m S_m(I_c + I_{sa} + I_a) - \mu_c S_m
\end{aligned} \quad (5)
$$

Note: $1/\delta_c = 1/(\delta_m + \omega_M)$ i.e., a calf that received maternal immunity will enter the susceptible calf class at the ($\delta_m$) calf ageing rate plus calf loss of immunity rate ($\omega_M$)

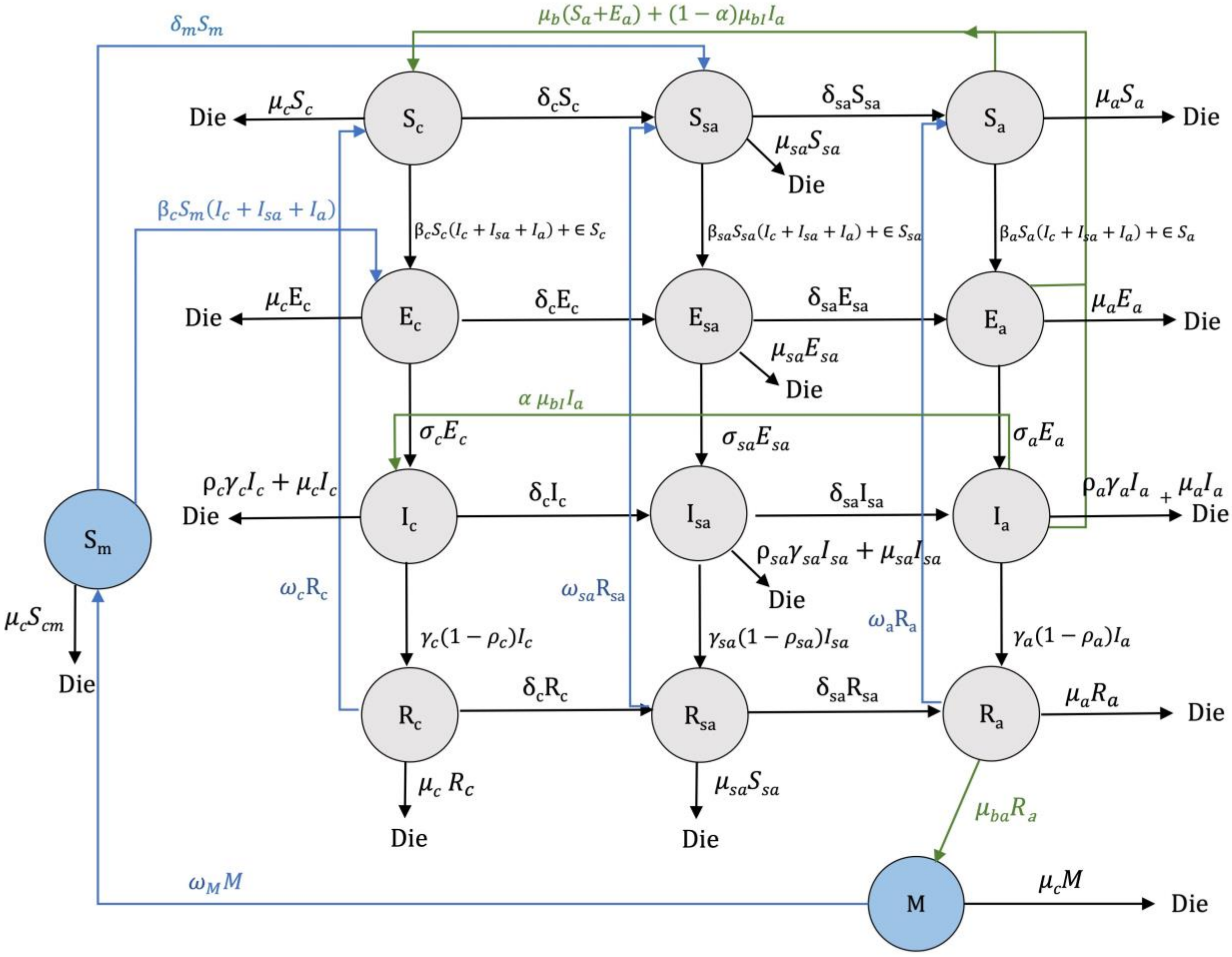


Figure S12 *SEIRMS/E* model of FMD and brucellosis.

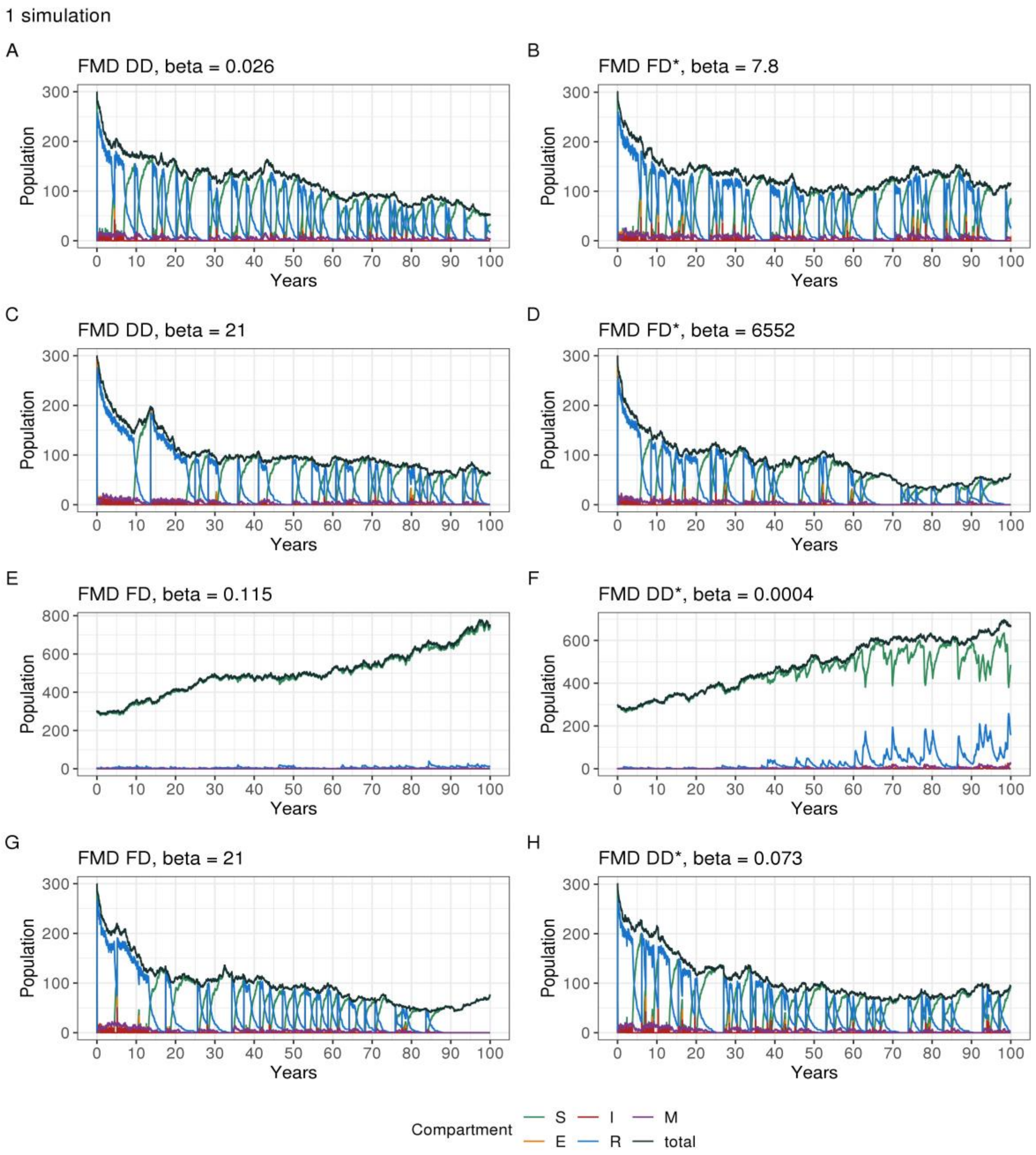


Figure S13 Single stochastic simulations of FMD. FMD FD with β = 0.115 (E) had the least population impact, while the other models decreased the total population. All models with different in β rates (0.026, 21) showed a cyclic pattern of outbreaks around every 5 years, except E & F.

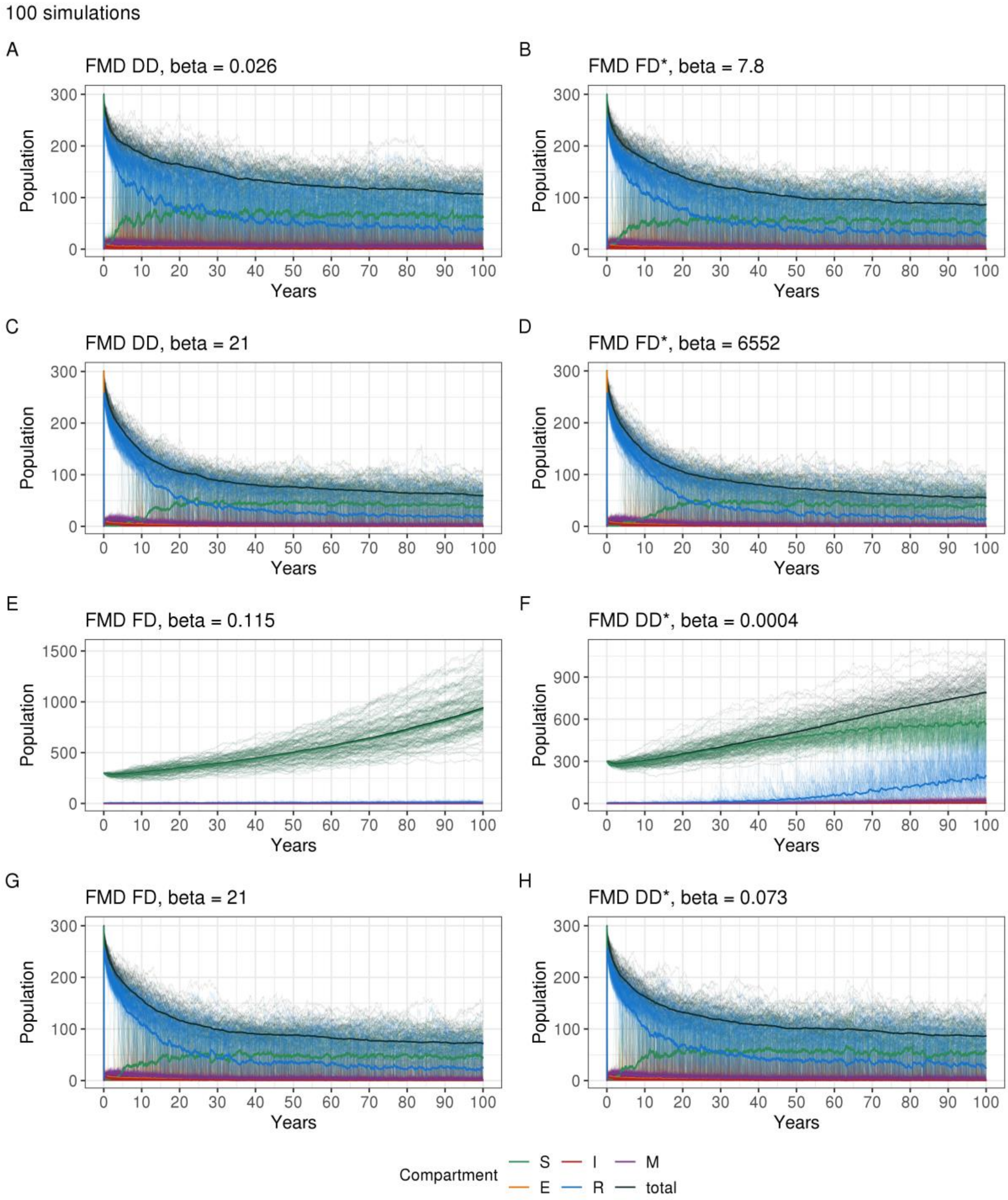


Figure S14 Hundred stochastic simulations of FMD.

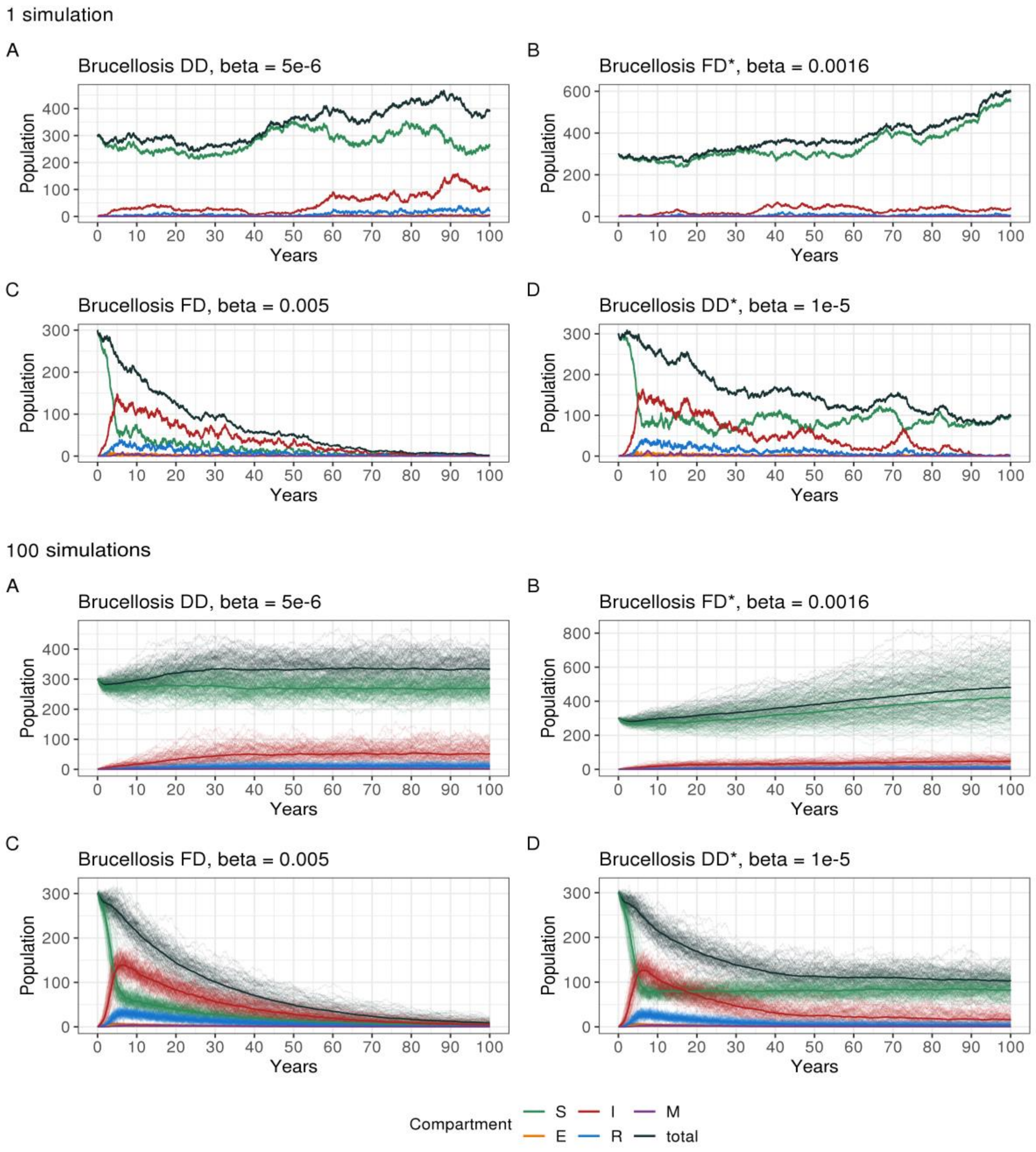


Figure S15 Single and 100 stochastic simulations of brucellosis. Brucellosis with FD transmission (C, D) had the most impact on populations, with a chance of population decline to zero. DD transmission showed a constant (A) and increase (B) in the populations.

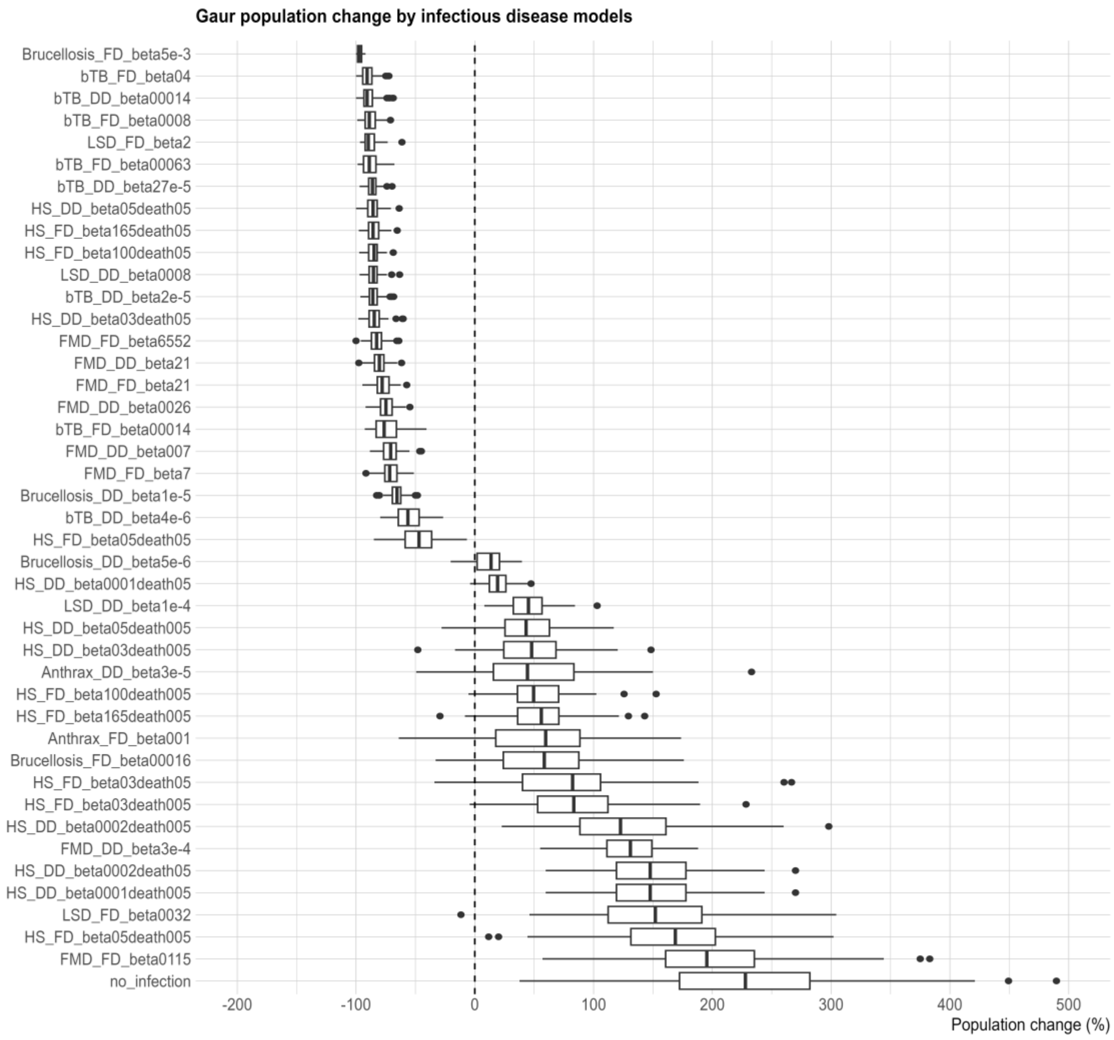


Figure S16 The average of the total population changes for all models. The boxplot presents the average percentage of the total population change compared to the no infection model (M1) to all the infectious disease models from the lowest to the highest (M2 to M43) of the population change in model simulations (N=100).

Table S1 The average of the total population changes for all models.

| No. | Model name | Model code | Average of the total population changes (%) |
|---|---|---|---|
| 1 | no_infection | M1 | 228 |
| 2 | FMD_FD_beta0115 | M2 | 200 |
| 3 | HS_FD_beta05death005 | M3 | 167 |
| 4 | LSD_FD_beta0032 | M4 | 155 |
| 5 | HS_DD_beta0001death005 | M5 | 149 |
| 6 | HS_DD_beta0002death05 | M6 | 149 |
| 7 | FMD_DD_beta3e-4 | M7 | 130 |
| 8 | HS_DD_beta0002death005 | M8 | 127 |
| 9 | HS_FD_beta03death005 | M9 | 85 |
| 10 | HS_FD_beta03death05 | M10 | 79 |
| 11 | Brucellosis_FD_beta00016 | M11 | 60 |
| 12 | Anthrax_FD_beta001 | M12 | 57 |
| 13 | HS_FD_beta165death005 | M13 | 55 |
| 14 | HS_FD_beta100death005 | M14 | 53 |
| 15 | Anthrax_DD_beta3e-5 | M15 | 51 |
| 16 | HS_DD_beta03death005 | M16 | 47 |
| 17 | HS_DD_beta05death005 | M17 | 46 |
| 18 | LSD_DD_beta1e-4 | M18 | 45 |
| 19 | HS_DD_beta0001death05 | M19 | 20 |
| 20 | Brucellosis_DD_beta5e-6 | M20 | 11 |
| 21 | HS_FD_beta05death05 | M21 | -48 |
| 22 | bTB_DD_beta4e-6 | M22 | -55 |
| 23 | Brucellosis_DD_beta1e-5 | M23 | -66 |
| 24 | FMD_FD_beta7 | M24 | -71 |
| 25 | FMD_DD_beta007 | M25 | -71 |
| 26 | bTB_FD_beta00014 | M26 | -74 |
| 27 | FMD_DD_beta0026 | M27 | -74 |
| 28 | FMD_FD_beta21 | M28 | -78 |
| 29 | FMD_DD_beta21 | M29 | -80 |
| 30 | FMD_FD_beta6552 | M30 | -83 |
| 31 | HS_DD_beta03death05 | M31 | -84 |
| 32 | bTB_DD_beta2e-5 | M32 | -85 |
| 33 | LSD_DD_beta0008 | M33 | -85 |
| 34 | HS_FD_beta100death05 | M34 | -85 |
| 35 | HS_FD_beta165death05 | M35 | -85 |
| 36 | HS_DD_beta05death05 | M36 | -86 |
| 37 | bTB_DD_beta27e-5 | M37 | -86 |
| 38 | bTB_FD_beta00063 | M38 | -88 |
| 39 | LSD_FD_beta2 | M39 | -88 |
| 40 | bTB_FD_beta0008 | M40 | -88 |
| 41 | bTB_DD_beta00014 | M41 | -89 |
| 42 | bTB_FD_beta04 | M42 | -90 |
| 43 | Brucellosis_FD_beta5e-3 | M43 | -97 |

**PCA biplot**

We conducted PCA using four disease parameters: beta transmission rate, incubation period, infectious period, and fatality rate. PCA can identify the contribution of these parameters to the population change. Also, PCA can show which disease has the similar traits grouping by the disease parameter and grading the impact based on the percentage of the population change from high (red) to low impact (blue). The beta transmission rate and incubation period demonstrated the highest contribution to the FMD FD model (rescaling DD, $\beta$ = 6552). Nevertheless, the fatality rate had the most significant impact on Anthrax FD and its DD (rescaling FD) with a 100% fatality rate but less effect on the population change. The other diseases showed less influence by a single parameter but had more impact on the population change (population decrease, indicated in red in Figure S17 – Figure S18), including bTB and Brucellosis.

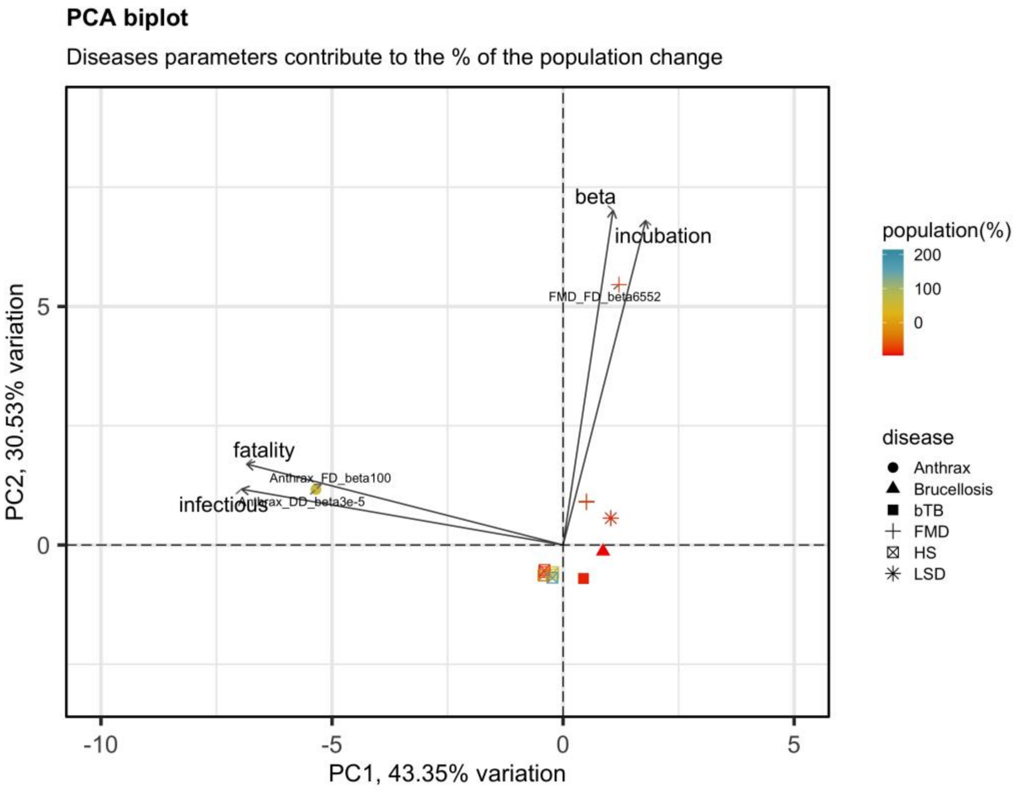


Figure S17 PCA biplot of 4 disease parameters, including beta transmission rate, incubation period, infectious period and fatality rate, contributing to the percentage of the total population change.

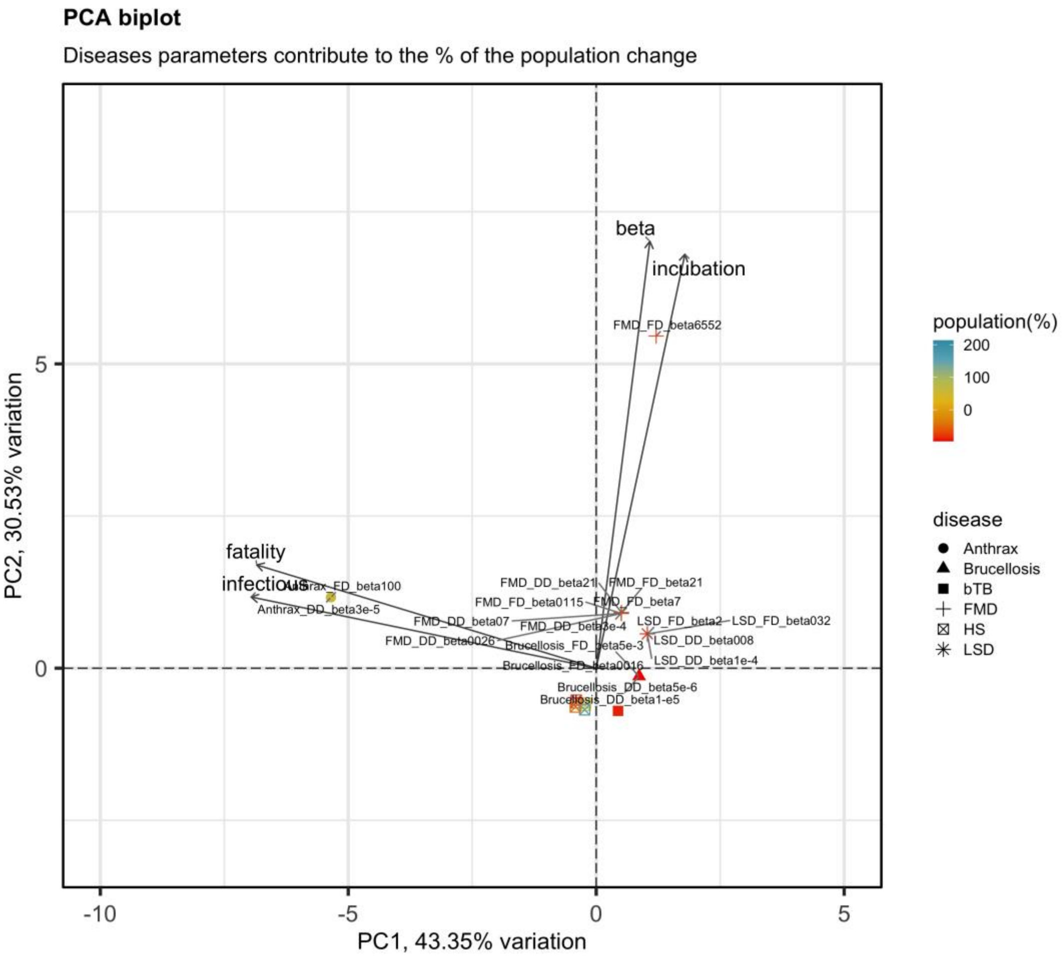


Figure S18 PCA biplot of 4 disease parameters with model's name, including beta transmission rate, incubation period, infectious period and fatality rate, contributing to the percentage of the total population change.
